# Protozoal populations drive system-wide variation in the rumen microbiome

**DOI:** 10.1101/2024.12.05.626740

**Authors:** Carl M. Kobel, Andy Leu, Arturo Vera-Ponce de León, Ove Øyås, Wanxin Lai, Ianina Altshuler, Live H. Hagen, Rasmus D. Wollenberg, Mads T. Søndergaard, Cassie R. Bakshani, William G. T. Willats, Laura Nicoll, Simon J. McIlroy, Torgeir R. Hvidsten, Oliver Schmidt, Chris Greening, Gene W. Tyson, Rainer Roehe, Velma T. E. Aho, Phillip B. Pope

## Abstract

While rapid progress has been made to characterize the bacterial and archaeal populations of the rumen microbiome, insight into how they interact with keystone protozoal species remains elusive. Here, we reveal two distinct rumen community types (RCT-A and RCT-B) that are not strongly associated with host phenotype nor genotype but instead linked to protozoal community patterns. We leveraged a series of multi-omic datasets to show that the dominant *Epidinium* spp. in animals with RCT-B employ a plethora of fiber-degrading enzymes that present enriched *Prevotella* spp. a favorable carbon landscape to forage upon. Conversely, animals with RCT-A, dominated by genera *Isotricha* and *Entodinium*, harbor a more even distribution of fiber, protein, and amino acid metabolizers, reflected by higher detection of metabolites from both protozoal and bacterial activity. We reveal microbiome variation across key protozoal and bacterial populations is interlinked, which should act as an important consideration for future development of microbiome-based technologies.

## Introduction

The herbivore rumen is a highly specialized organ that has co-evolved in symbiosis with a complex microbiome, made up of thousands of microbial populations whose interactions collectively convert plant material into energy-yielding metabolites for the host. The rumen microbiome acts as an interface between the nutrient potential of the feed and the metabolism of the host animal, and includes members from all domains of life: Archaea, Bacteria, and Eukarya (ciliate protozoa and fungi)^1,2^. From ingested plant material, cellulose, pectin, xylans, xyloglucans, and other polysaccharides are degraded by microbially encoded carbohydrate-active enzymes (CAZymes) down to their component monosaccharide units, which are subsequently fermented into several intermediates. Most importantly, pyruvate is converted to volatile fatty acids (VFAs) such as acetate, propionate, and butyrate^3^. Along this fermentation pathway, hydrogen (H_2_) is produced, which predominantly flows into methanogenesis but can also be incorporated into VFAs through alternative hydrogen sinks such as the reduction of fumarate^4^. The rumen epithelial wall is able to transport most of the VFAs directly into the blood, whereas more complex metabolites take a longer path, being assimilated by the posterior gastrointestinal tract^5^.

Rumen microbiome structure and function is shaped by many dynamic host-associated variables, such as diet, age, health status, animal husbandry, behavior, and breed. Efforts to monitor and predict overall rumen microbiome function for the purpose of improved animal production have mainly focused on recovering isolates and genomes of the various microbial populations. However, the superior amenability of bacteria and archaea to current molecular microbiology techniques has created significant domain-specific information bias, with recovery of greater than 50,000 bacterial and archaeal genomes compared to ∼50 for eukaryotic species^1,2^. The ciliate protozoa, specifically the class Litostomatea, subclass Trichostomatia, have a relatively large biomass in the rumen (up to 50%^1^), and are ubiquitous among ruminants. Although single-celled, they have complex organelles and physiological features, such as mouthlike adoral openings that lead to a tongue-like extrusible peristome, which ingests feed particles into an esophagus-like structure^6^. This, combined with their outside being covered with undulating cilia for propulsion, makes many of them voracious predators^6^. To add to their versatility, they express carbohydrate-active enzymes (CAZymes) and are able to degrade plant fibers^2^. Decades of in vitro work have shown that rumen ciliates often act as a microhabitat for archaea and bacteria^7^, especially *Methanobrevibacter* spp., which form metabolic mutualisms with several ciliate species by recycling the H_2_ produced by the ciliates as a main metabolic end product^8,9^. Providing *in vivo* context to the wider ecological impacts of rumen protozoal populations has proven immensely challenging but is necessary to advance microbiome-based solutions to animal productivity and sustainability, for example in the context of methane mitigation and improving feed efficiency.

Rapid advancement of biotechnological tools has improved the availability of data for resident rumen microbiota, yet information on how species interact within these multidomain ecosystems is still limited. In this study, we applied long-read metagenomics, existing single-cell amplified eukaryotic genomes, and genome-centric multi-omics of both host and its microbiome to improve resolution of inter-domain relationships and the influence they exert at a system level. Two breeds of cattle from a highly controlled experiment were phenotyped for key performance traits, and rumen contents, epithelial, and liver samples were analyzed across all molecular layers—genes, transcripts, proteins, and metabolites (**Fig. 1**). Taxonomic analysis identified two clear rumen microbiome structure types across the entire animal cohort that were not strongly correlated to breed, any of the recorded animal performance metrics, or methane emissions. Deeper analyses across the microbial domains identified two distinct protozoal population types that we hypothesize to drive system-wide microbiome differences, ultimately affecting the interlinked metabolisms that channel the flow of nutrients across the feed-microbiome-host axis.

**Figure 1.**
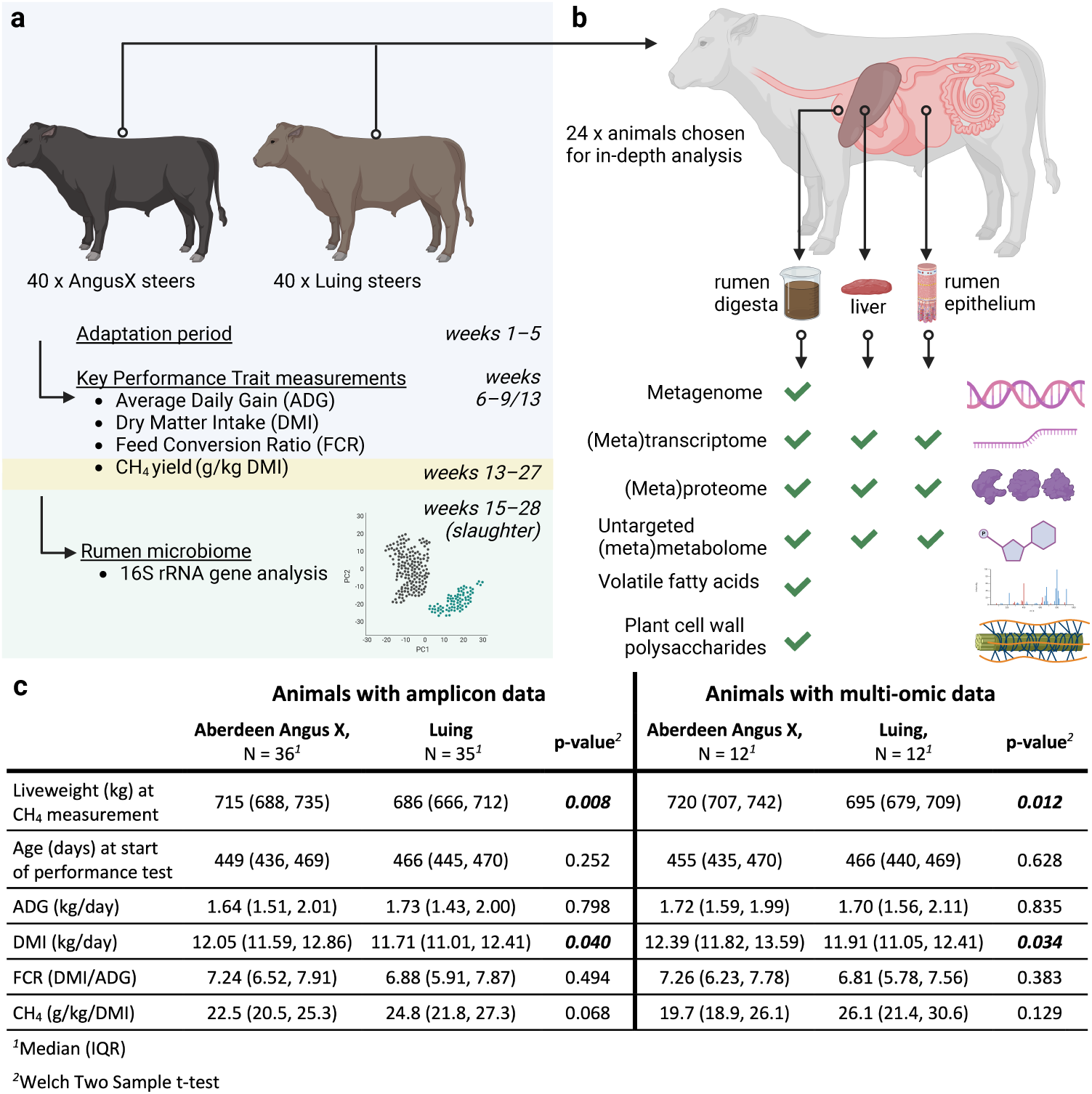
Experimental design of the animal trial and microbiome analyses. **a.** Animal trial setup. A total of 80 animals across two breeds (Aberdeen-Angus cross and Luing) were enrolled of which 71 completed the 3-6 month study period that culminated in their slaughter. Key performance traits such as average daily gain (ADG), dry matter intake (DMI), feed conversion ratio (FCR) and methane yield (g/kg DMI) were measured for all animals and rumen samples periodically collected across the duration of the trial. **b.** Sampling design for a subset of 24 animals, selected on the widest recorded level of natural methane yield variation. At slaughter, three sample locations were collected: Rumen digesta, rumen wall tissue, liver. Samples were characterized on several molecular layers: Genomes, transcripts, proteins, untargeted metabolomics. **c.** Key performance traits and other animal production metrics that were determined for all enrolled animals. IQR: interquartile range. Significant p-values are marked with bold italics.

## Results

### One controlled animal experiment reveals two distinct rumen microbiome structure types

As part of an effort to improve the depth of understanding within the rumen microbiome, we analyzed samples from a controlled feedlot trial of adult beef cattle fed a total mixed ration of forage and concentrate (ratio: 51:49). From an initial 80 animals representing two breeds that commenced the trial, 36 Aberdeen-Angus cross (AAX) and 35 Luing animals completed the experimental period with all planned measurements, including key performance traits (KPTs) such as dry matter intake (DMI), average daily gain (ADG), feed conversion ratio (FCR) and methane yield (g/kg DMI). For microbiome analysis, rumen samples were taken for all 71 animals at five timepoints across the experimental period and subjected to 16S rRNA gene amplicon sequencing, with an additional final time point sampled at slaughter (**Fig. 1a**). A subset of 24 animals (12 AAX, 12 Luing), representing the highest and lowest natural levels of methane yield, were sampled across both the host and its microbiome at slaughter. The datasets generated from these 24 animals included long-read metagenomics for metagenome-assembled genome (MAG) reconstruction as well as RNA, protein and metabolite analysis of rumen digesta, rumen epithelia and liver tissue (**Fig. 1b**). As expected, the recorded KPTs showed breed-dependent differences in animal metrics, such as a higher liveweight and dry matter intake (DMI) in AAX animals, and a trend for higher methane emissions (g/kg DMI) in Luing animals (**Fig. 1c**).

For microbiome characterization, long-read metagenomic sequencing of rumen samples from the 24-animal subset produced a total of 700 high- and medium-quality dereplicated metagenome-assembled genomes (MAGs, 656 classified as bacterial, 44 as archaeal; **Supplementary Table 1a**). These sample-specific MAGs, together with previously published fungal genomes (n=9)^10^ and protozoal single amplified genomes (SAGs) (n=53)^2,11^, formed the reference database for metatranscriptomic and metaproteomic analyses. Rumen metatranscriptomics identified 1,669,849 expressed genes (1,299,827 from bacteria, 80,325 from archaea, 252,768 from protozoa, and 9,529 from fungi), whereas metaproteomics identified 35,655 protein groups (16,823 from bacteria, 380 from archaea, 18,000 from protozoa, 137 from fungi, and 315 from the cattle host) (**Supplementary Table 1b**). To further assist our interpretations of host and microbial metabolic activity we generated untargeted metabolomic data from the three different sample types available (numbers of identified metabolites: rumen: 496; rumen epithelium: 517; liver: 859; **Supplementary Table 1b**). Finally, we performed VFA measurements from rumen fluid, as well as Microarray Polymer Profiling (MAPP) of rumen digesta, determining the composition and relative abundance of glycans available to the rumen microbiome.

Microbiome analysis of the 71 animals using the 16S rRNA gene sequence data did not reveal clear associations for any alpha or beta diversity metrics with breed, methane yield, or any of the other measured animal KPT (**Extended Data Fig. 1**). However, beta diversity plots illustrated two groups of animals whose microbiome structure distinctly clustered together, which could also be captured using probabilistic modeling (Dirichlet Multinomial Mixtures^12^) (**Fig. 2a**). Surprisingly, these two clusters, hereafter referred to as Rumen Community Type-A and -B (RCT-A and RCT-B), did not correspond to any measured animal KPT nor to any technical grouping that arose from the experimental workflow (**Extended Data Fig. 2a**). Furthermore, these community types were stable across time: the animals consistently stayed in the same cluster over the six timepoints sampled during the experiment (**Extended Data Fig. 2b**). Using genome-scale metabolic models to predict the metabolic reaction abundances of archaea and bacteria, we found little variation in functional potential between RCT-A and RCT-B, supporting the importance of including all microbial domains (e.g. eukaryotes) as well as functional omics beyond genes and genomes in our analyses (**Extended Data Fig. 3**).

**Figure 2.**
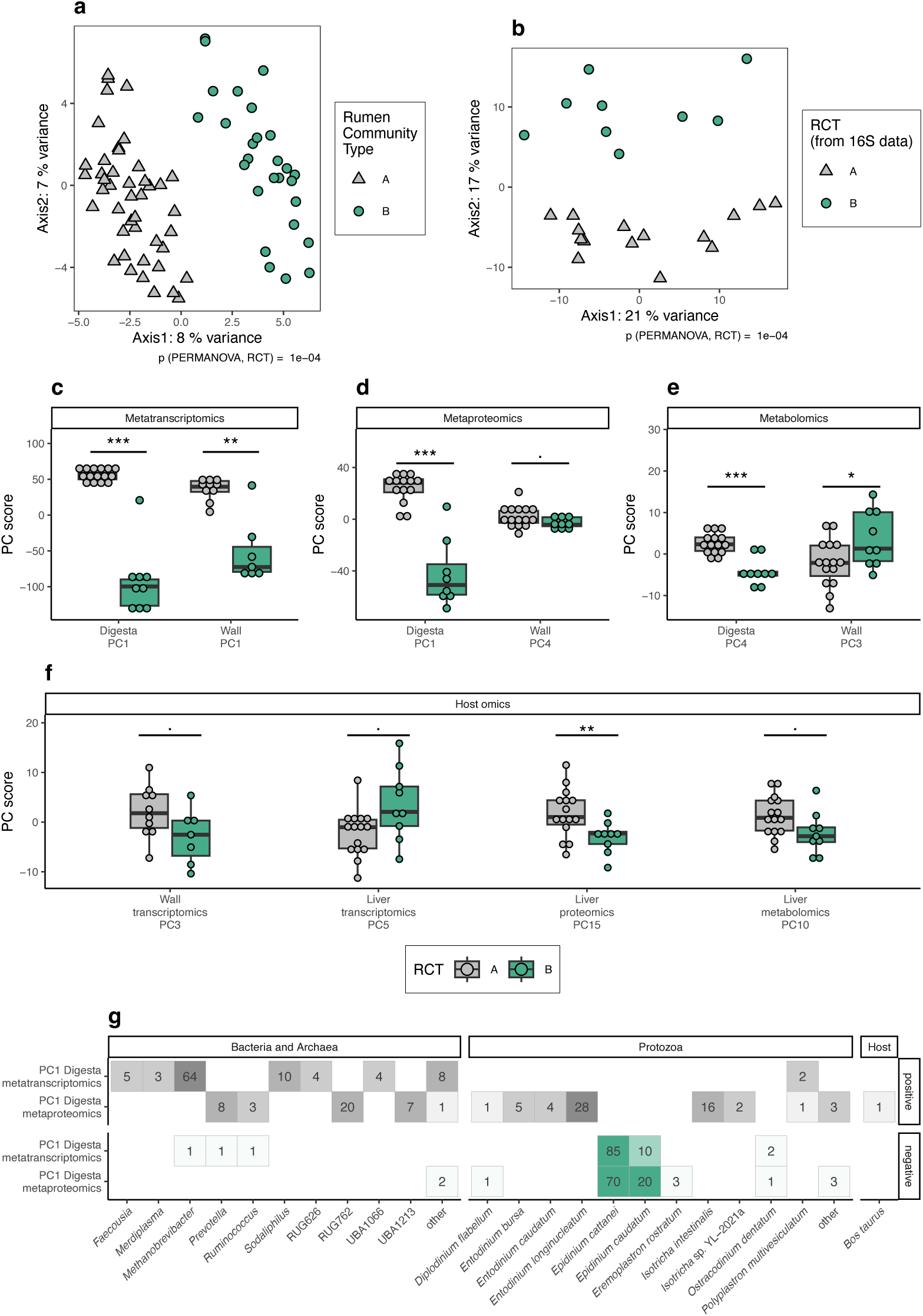
Rumen microbiome analyses showing two distinct groups, labeled as Rumen Community Type-A and -B. **a.** Principal Coordinate Analysis (PCoA) plot of robust Aitchison distances of 16S rRNA gene amplicon sequence analysis of 71 animals across two breeds (AAX = 36, Luing = 35), showing the two optimal clusters based on Dirichlet Multinomial Mixtures. **b.** PCoA plot of robust Aitchison distances of shotgun metagenomic data from a subset of 24 animals with the highest and lowest methane yield (g/kg DMI). **c-f.** Dot and box plots of microbiome-derived **c.** metatranscriptomic, **d.** metaproteomic, and **e.** metabolomic data as well as **f.** host-omic data, showing the first Principal Component (PC) with a significant difference between the two clusters (t-test) for each sample material and measurement type. Box hinges represent 1st and 3rd quartiles, and whiskers range from hinge to the highest and lowest values within 1.5*IQR of the hinge. **g.** Taxonomic classifications of features (transcripts or proteins) with the strongest contributions (i.e. loadings) to PC1 in digesta metatranscriptomics and proteomics, grouped based on association with rumen community type (positive: RCT-A; negative: RCT-B), showing the top 100 features for each omic data type and RCT; the "other" categories summarize taxa with only one transcript or protein in the top 100.

Curiously, RCT-A and -B were detectable across several omic layers in the 24-animal subset that we analyzed in more detail. Principal coordinates analysis (PCoA) of MAG abundances reflected the same pattern that was detected in the 16S rRNA gene sequence data (**Fig. 2b**). In Principal Component Analyses (PCA) of digesta and rumen wall epithelium metatranscriptomics as well as digesta metaproteomics, the first principal components (PCs) clearly differentiated RCT-A and -B, thus mirroring the 16S rRNA gene sequence analysis results of the entire animal cohort (**Fig. 2c-d**). The congruence between the molecular layers affirmed that elements of metabolism are affected by this distinct compositional difference in microbial communities of RCT-A and RCT-B. Untargeted metabolomics of digesta and the rumen wall epithelium also reflected this pattern on PCs 4 and 3, respectively (**Fig. 2e**). Finally, host proteomics and transcriptomics from wall and liver data showed a trend towards the two community types, although not always statistically significant (p < 0.05 only for PC15 from liver proteomics; 0.1 > p > 0.05 for other host data; **Fig. 2f**).

### Protozoal patterns associate with rumen community types

To further investigate the defined RCT-A or -B types, we extended our analyses to include domains of life present within the rumen samples of this study, incorporating the archaeal and bacterial MAGs as well as the single amplified genomes (SAGs) for protozoal populations. The taxonomic classifications of the transcripts and proteins with the strongest contributions (i.e. loadings) to the significant principal component patterns that were observed (**Fig. 2g**) clearly indicated that the RCT-A and -B clustering extended to the abundance profiles of detected protozoal species. Based on these and the differential abundance comparisons of metatranscriptomic and metaproteomic data, animals that exhibited the RCT-A pattern were enriched with families Entodiniinae and Isotrichidae, including high abundances of *Entodinium bursa*, *Entodinium caudatum*, *Entodinium longinucleatum*, and *Isotricha intestinalis*, as well as *Ostracodinium gracile* and *Polyplastron multivesiculatum* (**Fig. 3**). Conversely, animals with the RCT-B pattern were enriched with subfamilies Diplodiniinae and Ophryoscolecinae^2^ and typically included the species *Diplodinium dentatum, Epidinum cattanei*, *Epidinium caudatum,* and *Ophryoscolex caudatus* (**Fig. 3**).

**Figure 3.**
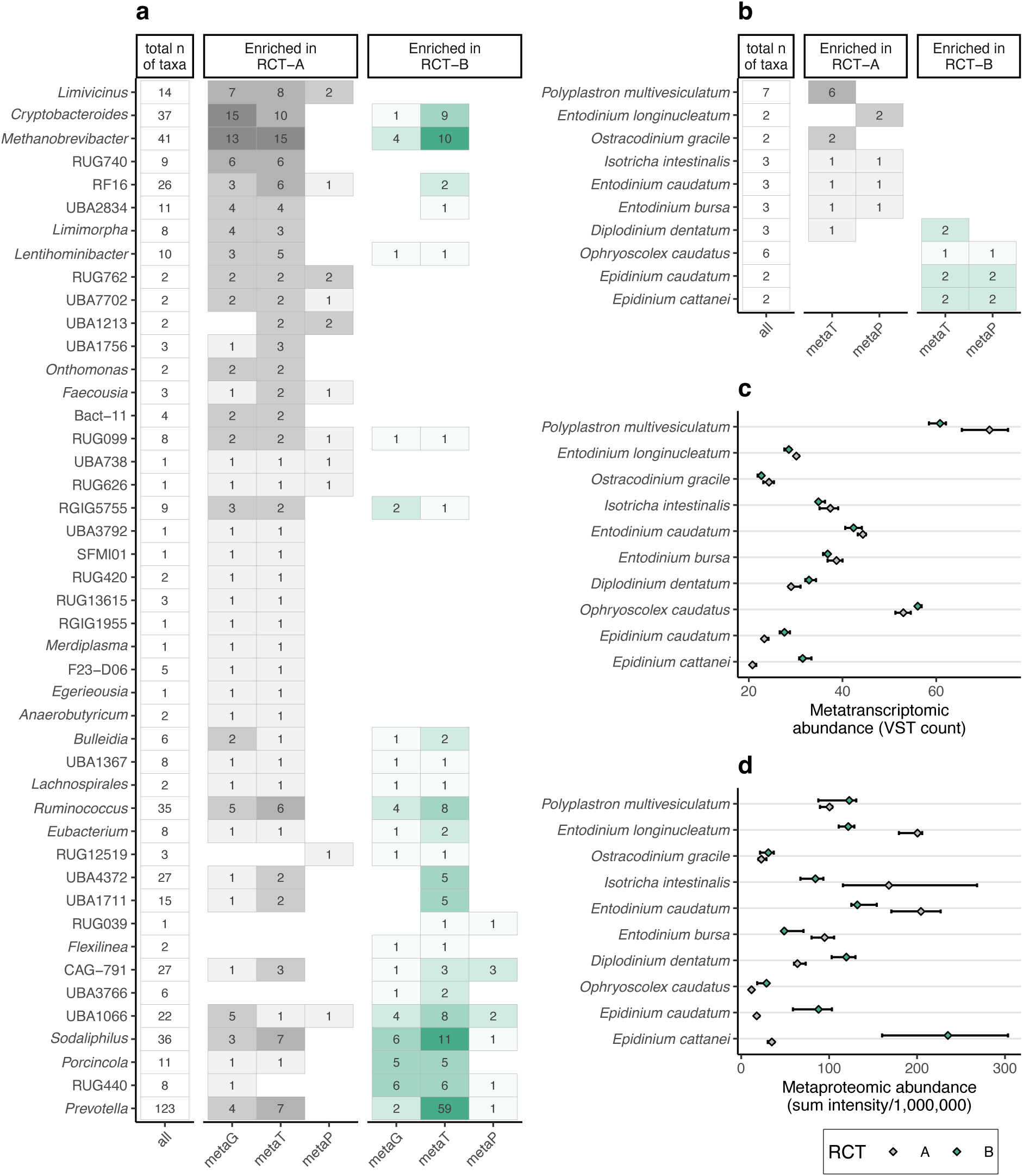
Differential abundances of taxa across omic approaches. a.-b. Summary of differential abundance results. Numbers indicate significantly different taxa (adjusted p < 0.05) out of the total indicated in the “Total n of taxa” column. metaG: taxon relative abundance compared with Wilcoxon Rank Sum tests; metaT: transcript counts summarized per taxon, compared with DESeq2; metaP: LFQ intensities summarized per taxon, compared with Wilcoxon Rank Sum tests. Multiple comparison correction is included in the testing procedure for DESeq2 (Benjamini-Hochberg FDR); additional FDR correction was implemented for Wilcoxon Rank Sum tests. **a.** Metagenome-assembled genomes (MAGs) per genus for bacteria and archaea, showing genera with differentially abundant MAGs supported by at least two omics in the same direction. **b.** Single-amplified genomes (SAGs) per species for ciliate protozoa, showing species with at least two differentially abundant results. **c.-d.** Abundances of the protozoal species from panel b. Diamonds indicate medians, whiskers IQR. **c.** Metatranscriptomic data, **d.** Metaproteomic data.

Coexistence or exclusion patterns of protozoal species in ruminants have been observed for over half a century, initially in microscopy-based studies suggesting that certain protozoal species are mutually exclusive, both in sheep and cattle^13^. These early, microscopy-based studies suggested that protozoal populations could be classified into two community types: type A, defined by the presence of *Polyplastron multivesiculatum*, often accompanied by *Diploplastron affine*, and type B, characterized by *Epidinium* and *Eudiplodinium* spp., together or alone^13^. The protozoal abundance patterns observed in our RNA and protein data bore remarkable similarities to these community types. A noticeable difference between the classical community types and our results was the coexistence of *P. multivesiculatum* and *Epidinium* spp. in our RCT-B animals. Such coexistence has been suggested to constitute an AB community type^7,139,14,15^, and postulated to represent a transitional stage from type B to A^9^. To explore the interrelationships of these protozoa, we examined the metatranscriptomes and metaproteomes of rumen samples collected from six animals over a period of three to six months. In line with the temporal stability of the bacterial and archaeal community structure (**Extended Data Fig. 2b**), our data indicated a constant, low but detectable presence of *P. multivesiculatum* together with *Epidinium* spp. in RCT-B animals (**Extended Data Fig. 4**). This brings to question whether these two protozoal genera are indeed mutually exclusive.

#### Protozoal community types affect bacterial and archaeal structure and function

We sought to better identify the microbial populations driving the observed system-wide variation, as well as its function implications, by further examining the metatranscriptome and metaproteome data. Differential expression analysis and the features with the strongest loadings in our abovementioned PCA analysis both highlighted that specific bacterial, archaeal and protozoal populations were indeed more prevalent in either RCT-A or -B (**Fig. 2g**, **Fig. 3**). Collectively, for animals categorized as RCT-A, the metaproteomes from their rumen were largely dominated by *Isotricha* spp, *Entodinium* spp, and the clostridial lineage *Acutalibacteraceae* (RUG762) while transcriptomes for various *Methanobrevibacter* spp., *Sodaliphilus* spp., *Faecousia* spp. and *Lachnospiraceae* (UBA1066) were also prevalent. In contrast, both the metatranscriptome and metaproteome for rumen samples from RCT-B animals showed far higher detection of *Epidinium* spp., while metatranscriptomics also indicated an enrichment in *Prevotella* spp. (**Fig. 3**).

To link biology to these observed structural patterns, we explored the annotated functions of the differentially detected populations more deeply with specific attention to the key functional stages of rumen digestion, namely fiber hydrolysis, fermentation of organic material and production of energy-yielding volatile fatty acids (**Fig. 4**). By far the most active fibrolytic population observed in RCT-B animals was *Epidinium* spp. which contained a plethora of CAZymes predicted to act upon cellulose, arabinoxylans, beta-mannans and arabinogalactan protein glycans commonly found in grasses and grains (**Fig. 4b**). Epidinia are the most reputable among the rumen ciliates to actively attach and degrade plant material, as visually confirmed across a series of prior studies^16^. In a scenario where epidinia are more proliferant and engaging in plant material deconstruction, it is reasonable to expect their activity and size will impact the glycan landscape that is available for neighboring microbial populations. Indeed, our MAPP analysis of rumen digesta particles was suggestive of differences in various beta-glucan, xylans, xyloglucans, and arabinogalactan proteins between the epidinia-dominated RCT-B animals and the RCT-A animals (**Fig. 5a**). In turn, many fiber-degrading bacterial lineages, such as *Sodaliphilus* and *Prevotella* spp., were detected at higher levels in metatranscriptomic data arising from RCT-B animals (**Fig. 3a**), supporting our hypothesis that system-wide effects are driven by protozoal activity. While metaproteomic detection of central butyrate-producing enzymes was observed in both RCT’s, we suspect the elevated activity of epidinia species was influential towards higher butyrate levels in RCT-B animals (**Fig. 4b-c**), which is supported by prior meta-analysis of protozoa that showed defaunation will substantially decrease ruminal butyrate levels^17^.

**Figure 4.**
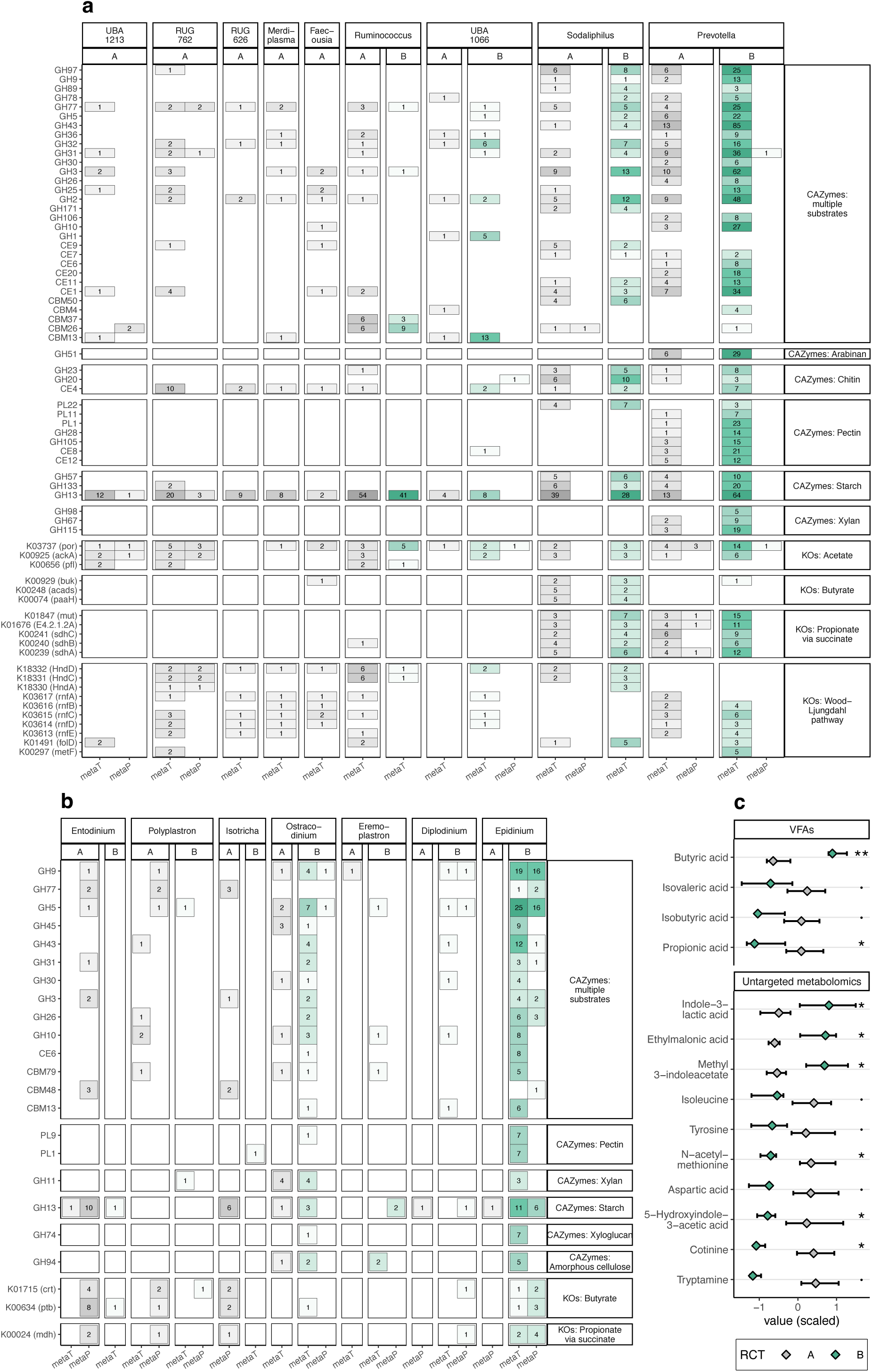
Functional differences between rumen community types. a.-b. Summaries of KEGG Orthologs in pathways of interest (selected based on Ni, G. et al. 2024) and CAZymes in **a.** bacterial and **b.** protozoal genera of interest. Genera are the same as listed in Fig. 2g. CAZymes and KOs were chosen by first selecting the 10 pathways or substrates (labels on right side of plot) with the largest amount of differentially abundant features between RCT-A and RCT-B, and then further trimming the list to those CAZymes and KOs (labels on left side of plot) that were detected as differentially abundant at least 5 times total, regardless of taxonomic origin or omic layer. metaT = metatranscriptomic data, showing numbers of transcripts upregulated in either RCT-A or RCT-B (DESeq2 adjusted p-value < 0.1). metaP = metaproteomic data, showing numbers of protein groups with higher intensities in either RCT-A or RCT-B (fdr-adjusted *p-*values from Wilcoxon Rank Sum test < 0.1). Complete pathways and associated KOs for each of the depicted genera are available in **Extended Data Fig. 6**. **c.** Summary of metabolite measurements, showing those volatile fatty acids (VFAs) or metabolites from untargeted metabolomics with annotation level 1 or 2a that had a multiple comparison corrected *p-*value < 0.1. Diamonds indicate medians, whiskers IQR. ** p < 0.01, * : p < 0.05, · : 0.1 > p > 0.05.

**Figure 5:**
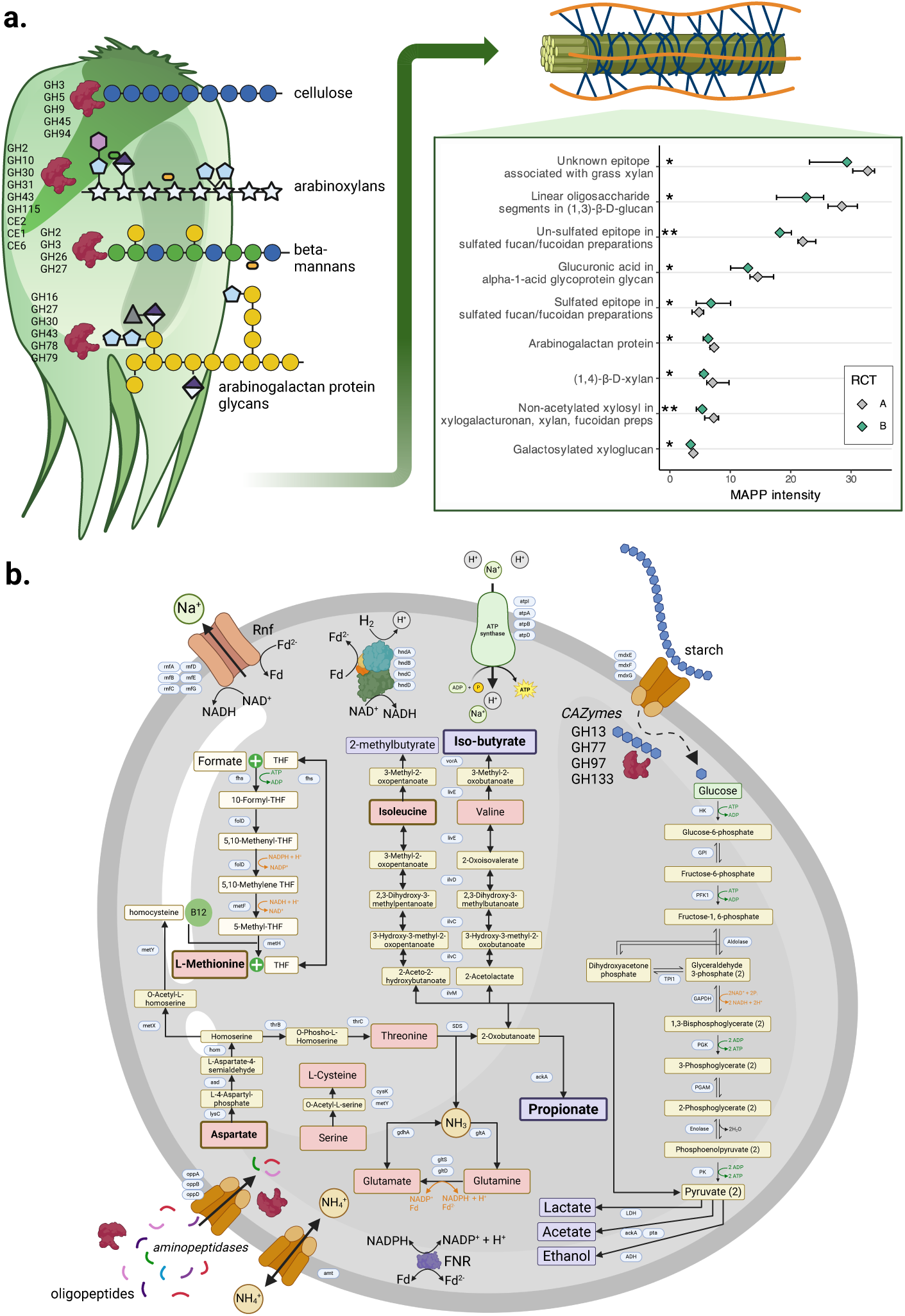
Metabolic predictions from major populations strongly featured in RCT-A and -B animals that are predicted to influence rumen function. **a.** Illustration of *Epidinium cattanei*, the protozoal species most strongly associated with RCT-B, predicted to engage a broad range of CAZymes to degrade dietary plant material. Given the size and activity of *E. cattanei* their fiber degrading metabolism is predicted to impact the rumen structure and function of RCT-B animals, which was supported via Microarray Polymer Profiling (MAPP) of various hemicellulosic plant fibers, which highlighted differential abundances. In the MAPP inset, diamonds indicate medians, whiskers interquartile ranges, and stars represent uncorrected *p-*values from Wilcoxon rank sum tests. **b.** Pathway reconstruction for the *Acutalibacteraceae*-affiliated RUG762 population, strongly associated with RCT-A, highlighting, sugar fermentation, amino acid (red boxes) metabolism and a partial Wood-Ljungdahl Pathway, which was supported by the associated energy conservation machinery such as the electron-bifurcating hydrogenase (HndABCD, [FeFe] group A), ferredoxin:NAD-oxidoreductase (Rnf) complex, and components of a FoF1 ATP synthase. Bold text indicates differentially abundant metabolites from Fig. 4.

In the absence of elevated epidinia metabolism within RCT-A animals, both PCA and differential abundance analyses indicated the primary responsibilities for digestion was shared more broadly across the protozoal species *Entodinium* spp. and *P. multivesiculatum* as well as bacteria affiliated to family *Acutalibacteraceae* or genera *Faecousia* and *Merdiplasma* (**Fig. 4a-b**). The *Isotricha* species that dominated RCT-A animals were, as expected^17^, not primarily degraders of plant material, though we suspect their influence still impacted heavily upon bacterial populations. For example, populations affiliated to RUG762 (*Acutalibacteraceae*), had some of the strongest loadings for RCT-A animals within the metaproteomic PCA analysis (**Fig. 2g**) and similar to *Isotricha* species were consistently enriched in RCT-A (**Fig. 3a**). Functional annotation suggested that RUG762 populations were engaged largely in protein and amino acid metabolism, and this was supported by metaproteomic and metabolomic enrichment of the enzymes and metabolites for aspartate, glutamine and branched chain amino acid metabolism in RCT-A animals (**Fig. 4c**, **Fig. 5b**). Fermentation end products were predicted to be acetate and possibly propionate and branched-chain volatile fatty acids, which were also detected at higher proportions in RCT-A animals (**Fig. 4c**). The protein and amino acids for ruminal metabolism could plausibly arise from the grain fraction of the animal’s diet (355 g/kg dry matter in the concentrate component). However, *Isotricha* spp. have been shown to excrete cellular nitrogen in the form of amino acids, principally alanine, proline, glutamic acid, and aspartic acid^16,18^. If such excretion of amino acids indeed occurs in RCT-A animals dominated by *Isotricha* spp. our observations of elevated RUG762 metabolism are plausibly connected, though we acknowledge this hypothesis must be tested in future experiments that examine cellular proximity and nutrient transfer between these populations.

It was interesting to note that for RCT-A animals a grouping of *Methanobrevibacter*-affiliated populations were detected at significantly higher abundance and/or with PCA loadings clearly associating them with RCT-A, despite there not being significant differences in measured methane yield across the two groups of animals (**Extended Data Fig. 2a**). The holotrich *Isotricha* species have been repeatedly shown^17^ to associate with different methanogenic populations than entodiniomorphids (e.g. epidinia), and our data also followed this trend with *Methanobrevibacter* populations in epidinia-dominated RCT-B animals seemingly of distinct strains compared to RCT-A (**Fig. 3a, Extended Data Fig. 5**). Functional examination of bacterial populations enriched in RCT-A animals (**Fig. 3a**) also identified several taxa, including *Faecousia* and *Merdiplasma* species^19^, that were predicted to encode multimeric electron-bifurcating [FeFe] group A hydrogenases (HndABCD) as well as selected features putatively associated with the Wood-Ljungdahl pathway (WLP) (**Fig. 4a, Extended Data Fig. 6**). The WLP potentially facilitates reductive acetogenesis and can act as an alternative hydrogen sink to methanogenesis. While reductive acetogens indeed co-exist with methanogens, under normal rumen conditions they are believed to be outcompeted energetically and thus are often observed in low abundance. However, in a methane-inhibited rumen with elevated hydrogen partial pressure, *Faecousia* and *Merdiplasma* species have been observed to flourish^20^. Indeed, aforementioned RUG762 populations were also suspected to encode a partial WLP as well as the associated energy conservation machinery such as the bidirectional electron-bifurcating hydrogenase (HndABCD), ferredoxin:NAD-oxidoreductase (Rnf) complex, and FoF1 ATP synthase (**Fig. 5b**). However, the absence of the acetyl-CoA synthase/carbon monoxide dehydrogenase complex that is required for the complete reduction of CO_2_ to acetate, leads us to speculate that RUG762 populations are instead producing methionine via a cobalamin-dependent 5-methyltetrahydrofolate– homocysteine methyltransferase. In the context of higher methanogenic features in RCT-A animals, the non-differential methane yield levels observed across the RCT-A and -B animal are fascinating and are likely arising from as-yet undefined hydrogen flow that influences differing methanogen strains and other hydrogenotrophs in the rumen.

### Implications for the host animal

Despite the distinct systems-wide microbiome shifts that were reconstructed for RCT-A and - B animals, we were surprised to observe only limited effects of these microbial community differences on host biology. Animal performance measurements (**Extended Data Fig. 2a**), microbial and host metabolomic data, and host expression data in gut epithelial and liver tissues showed only minor changes to a small number of features (**Fig. 2**). The clearest difference was the relative composition of several amino acids and VFAs, with propionate and branched chain volatile fatty acids higher in RCT-A animals, while butyrate levels were higher in RCT-B (**Fig. 4c**). Since VFAs are the major energy source for the host animal and are taken up directly through the rumen wall epithelium^3^, we further applied a series of network analyses using rumen and epithelial proteomic data to ascertain if underlying expression patterns were indeed evident between metabolically linked microbial and host pathways. From rumen metaproteomes, weighted gene correlation network analysis (WGCNA)^21^ identified a wide variety of co-expression modules (ME) that contained mixtures of protozoal, bacterial and archaeal proteins; many of these modules were, unsurprisingly, strongly correlated with the RCT variable (**Extended Data Fig. 7**). In the epithelial proteomics data, WGCNA identified only two co-expression modules, comprised largely of host proteins, that were correlated with the RCT groupings, none of which were enriched with proteins functionally inferred in VFA metabolism (**Extended Data Fig. 8**). Of note, interlinked patterns of rumen digesta (ME9 and ME13) and epithelial (ME1) modules were enriched in proteins annotated in cysteine and methionine metabolism and RUG762 populations suggesting possible metabolic interplay of amino acids, though this needs future testing for validation. The lack of striking host effects arising from microbiome differences between RCT-A and -B animals highlights the extraordinary plasticity and functional redundancy of the rumen microbiome.

## Discussion

Rumen protozoa are large and complex compared to their bacterial and archaeal neighbors and their presence and distribution within the livestock rumen has been heavily documented for well over 130 years^16^. Despite their long-standing history their impact across the total rumen ecosystem remains poorly understood at a molecular level due to technical restrictions that have impeded their study, and which have only recently been overcome with omics methodologies. Herein we were excited to link the molecular patterns and functional interpretations in our data to community types first postulated over 60 years ago via light microscopy^13^. When first describing protozoal community types in 1962 J. Margaret Eadie explicitly stated: *“It is concluded that inter-relationships of the type described may play an important role in determining the components of a particular rumen microfauna.”*^13^. We show that for the animals in this study, the system-wide rumen microbiome structure indeed extended beyond the protozoal components originally proposed in community types A and B to encompass bacterial and archaeal populations.

Advancing the original Eadie hypothesis, our multi-layered omic datasets offered plausible interpretations on how two independent modes of metabolic interactions are interlinked across the rumen microbiome of RCT-A and -B animals. Of particular note was the seemingly direct influence certain protozoal species (e.g. *Epidinium* spp) play at higher trophic levels such as fiber hydrolysis, which likely impacts fiber structural configuration and availability for bacterial fibrolytic populations. On the other hand, protozoal metabolism of *Isotricha* spp. was predicted to indirectly affect how nutrients enter the food chain via excretion of metabolites such as amino acids and hydrogen, which impacted the structure and function of intermediate fermenters. While this study goes some way into explaining the microbiome-wide effects that particular protozoa can exert, major questions regarding the origin of their structural configuration still remain. We speculate the original protozoal seeding took place via animal-animal contact likely during early life transition that started with mother-calf contact and gradually extended to other animals across the greater herd. Unfortunately, behavioral data prior to animal enrolment and pen groupings used in this animal trial were not recorded, though it was clear that grouping of RCT-A and -B animals together in randomized pens had no immediate nor long term influence upon microbiome structure.

In this study we show that the acceleration in genome recovery of protozoal populations and their supplementation into rumen microbiome databases has massively impacted our ability to estimate the transdomain microbial trophic cascades that convert complex plant material into energy-yielding nutrients for the host animal. Moving forward, several outstanding knowledge gaps need to be prioritized so that greater microbiome resolution can be routinely gained. Laboratory-based experiments that validate both proximity and metabolic interactions between protozoa, bacteria and archaea will lead to improved interpretations of how protozoa modulate rumen biology and formulate tools to potentially intervene where desired. Finally, more extensive surveys of increased animal numbers, varying diets, breeds and management practices will need to be analyzed at a depth comparative to the present study to ascertain the wider implications of protozoal-bacterial-archaeal interactions, and how that knowledge can be applied to improve microbiome modulation strategies that make meaningful impact.

## Methods

### Ethics statement

The animal experiment was conducted at the Beef and Sheep Research Centre of Scotland’s Rural College (6 miles south of Edinburgh, UK). The experiment was approved by the Animal Experiment Committee of SRUC and was conducted in accordance with the requirements of the UK Animals (Scientific Procedures) Act 1986.

### Experimental design and measurement of key performance traits

An initial group of 80 animals representing two breeds of beef cattle (Aberdeen-Angus cross (AAX, n = 40), and Luing (n = 40) was selected for the experiment. Of these, 71 (AAX: n = 36; Luing: n = 35) successfully completed the designed sampling scheme. All animals were provided a typical basal diet consisting of whole crop barley (300 g/kg DM), grass silage (200 g/kg DM), barley (355 g/kg DM), maize dark grains (120 g/kg DM), molasses (15 g/kg DM) and minerals (10 g/kg DM). For half of the animals, the experimental design originally involved supplementation with *Asparagopsis taxiformis* red algae vegetative tissue (thallus) at 0.3% of the organic mass (OM). *A. taxiformis* is a feed additive which has been shown to reduce methane emissions in past studies^22–25^. However, due to adverse effects observed in animals during the planned three-week seaweed adaptation phase, supplementation was terminated after just 15 days. All animals were given a further 5 weeks to adapt to basal feed before performance testing was carried out. Temporal 16S rRNA gene amplicon analysis of samples collected before and after the seaweed supplementation indicated no long-lasting microbiome effects (Extended Data Fig. 2c). Due to this delay, the heaviest 32 animals, balanced for breed, underwent a shorter performance test period of 4 weeks instead of the normal 8 weeks. During performance testing, daily feed intake was recorded using electronic feeders (HOKO, Insentec, Marknesse, The Netherlands). Twice weekly, duplicated samples of each diet component were collected to determine dry matter content and to calculated dry matter intake (DMI). Body weight of each animal was measured weekly to estimate average daily gain (ADG) using a linear regression model including time on test. Feed conversion ratio (FCR) was calculated for each animal as average daily DMI divided by ADG.

At the end of the experimental period, the animals’ methane emissions were measured in respiration chambers. One week prior to entering the respiration chambers, the cattle were single-housed in training pens, identical in size and shape to the pens inside the chamber, to adapt to individual housing. The cattle were allocated to six respiration chambers based on the criterion of minimization of the variation in body weight. They remained in the respiration chambers for 3 days, which included one day for adaptation and a 48-hours measurement period for methane emissions.

Of the 71 animals that completed the trial, 24 were selected for multi-omic analysis, including equal numbers of the two breeds, and representing the full range of methane emissions. For a further subset of six animals (out of 24), samples were also analyzed for a time series collected during the experimental period using orogastric tubing, as described below.

### Rumen content and tissue sample collection

On live animals, longitudinal rumen fluid samples (50 ml) were collected using a stomach tube (16×2700 mm Equivet Stomach Tube; Jørgen Kruuse A/S, Langeskov, Denmark) nasally and aspirating manually. Samples were collected prior to the adaptation phase to seaweed, before and after the performance test as well as immediately after leaving the respiration chambers. Additionally, rumen fluid samples (50 ml) were obtained after the animals were slaughtered in a commercial abattoir, immediately after the rumen was opened to be drained. Immediately after sampling, the rumen digesta was filtered through two layers of muslin and a 5 ml sample of the filtered liquid was transferred into a 30ml universal containing 10 ml of PBS-Glycerol, then stored in a freezer at -80°C.

Rumen cell wall samples were collected from the central region of the ventral sac before the rumen had been washed. The ruminal tissue was dipped into a 125 ml beaker containing a PBS solution to remove the ruminal digesta. The tissue was sliced using a sterile scalpel and transferred to a 30 ml universal tube containing 5 ml RNALater. Additionally, liver samples were taken by the meat inspector, with a section cut out using a sterile scalpel and then stored in a 30 ml universal tube with 5 ml RNALater. All tissue samples were stored in a freezer at - 80 °C before being analyzed. Further details regarding the sampling and experimental procedures carried out at SRUC can be found in previously published studies^26,27^ which followed a similar protocol.

### 16S rRNA gene amplicon sequence data

Rumen digesta sample DNA extraction, PCR amplification and sequencing of 16S rRNA gene amplicons was performed at DNASense ApS (Aalborg, Denmark).

#### Sample DNA extraction

Rumen digesta DNA was extracted using the FastDNA Spin kit for Soil (MP Biomedicals, USA) with the following exceptions to the standard protocol: 500 μL of sample, 480 μL Sodium Phosphate Buffer and 120 μL MT Buffer were added to a Lysing Matrix E tube. Bead beating was performed at 6 m/s for 4×40s. Gel electrophoresis using Tapestation 2200 and Genomic DNA screentape (Agilent, USA) was used to validate product size and purity of a subset of DNA extracts. DNA concentration was measured using Qubit dsDNA HS/BR Assay kit (Thermo Fisher Scientific, USA).

#### Sequencing library preparation

Amplicon libraries for the 16S rRNA gene variable region 4 (abV4-C) were prepared using a custom protocol based on an Illumina protocol^28^. Up to 10 ng of extracted DNA was used for PCR amplification. Each reaction (25 μL) contained (12.5 μL) PCRBIO Ultra mix and 400 nM of each forward and reverse tailed primer mix. The PCR program was as follows: initial denaturation at 95 °C for 2 min, 30 cycles of amplification (95 °C for 15 s, 55 °C for 15 s, 72 °C for 50 s) and a final elongation at 72 °C for 5 min. Duplicate reactions were performed for each sample and the duplicates pooled afterwards. The primers targeting the abV4-C region were the following, designed according to^28^ : [515FB] GTGYCAGCMGCCGCGGTAA and [806RB] GGACTACNVGGGTWTCTAAT^29^, with tails that enable attachment of Illumina Nextera adaptors necessary for sequencing in a subsequent round of PCR. The amplicon libraries were purified using the standard protocol for CleanNGS SPRI beads (CleanNA, NL) with a bead to sample ratio of 4:5. DNA was eluted in 25 μL of nuclease free water (Qiagen, Germany). DNA concentration was measured using Qubit dsDNA HS Assay kit (Thermo Fisher Scientific, USA). Gel electrophoresis using Tapestation 2200 and D1000/High sensitivity D1000 screentape (Agilent, USA) was used to validate product size and purity of a subset of libraries.

Sequencing libraries were prepared from purified amplicon libraries using a second PCR. Each reaction (25 μL) contained PCRBIO HiFi buffer (1x), PCRBIO HiFi Polymerase (1 U/reaction) (PCRBiosystems, UK), adaptor mix (400 nM of each forward and reverse) and up to 10 ng of amplicon library template. PCR was done with the following program: initial denaturation at 95 °C for 2 min, 8 cycles of amplification (95 °C for 20 s, 55 °C for 30 s, 72 °C for 60 s) and a final elongation at 72 °C for 5 min. The resulting libraries were purified following the same protocol as above for the first PCR.

#### DNA sequencing

The purified sequencing libraries were pooled in equimolar concentrations and diluted to 2 nM. The samples were paired-end sequenced (2×300 bp) on a MiSeq (Illumina, USA) using a MiSeq Reagent kit v3 (Illumina, USA) following the standard guidelines for preparing and loading samples on the MiSeq. > 10 % PhiX control library was spiked in to overcome low complexity issues often observed with amplicon samples.

#### Sequence data analysis

Quality trimming and amplicon sequence variant (ASV) inference for the 16S rRNA gene amplicon sequence data was performed with dada2^30^ following the recommended Big Data Paired-end workflow^31^ using default parameters, except for the following choices for the filterAndTrim step: truncLen = 240 for forward, 200 for reverse reads; trimLeft = 20 for forward, 30 for reverse reads; maxEE = 2, and truncQ = 6. The reference database for taxonomic classification was the dada2 formatted version of release 214 of the Genome Taxonomy Database (GTDB)^32^.

### Metagenomics

DNA extraction and sequencing as well as initial metagenomic sequence data analysis for rumen digesta samples was performed at DNASense ApS (Aalborg, Denmark).

#### DNA extraction

DNA intended for sequencing on the Illumina platform was extracted during the workflow for 16S rRNA gene amplicon data, as described above. DNA intended for ONT sequencing was extracted with the DNeasy PowerSoil Kit (Qiagen, Germany) and further cleaned with the DNeasy PowerClean Pro Cleanup Kit (Qiagen, Germany). A custom SPRI (Solid Phase Reversible Immobilization) short fragment removal step was implemented to remove fragments shorter than approximately 1500-2000 bp. DNA concentration and purity was assessed using the Qubit dsDNA HS Assay kit (Thermo Fisher Scientific, USA) and the NanoDrop One device (Thermo Fisher Scientific, USA). DNA size distribution was evaluated using the Genomic DNA ScreenTape on the Agilent Tapestation 2200 (Agilent, USA).

#### Illumina sequencing

Extracted DNA was fragmented to approximately 550 bp using a Covaris M220 with microTUBE AFA Fiber screw tubes and the settings: Duty Factor 10 %, Peak/Displayed Power 75W, cycles/burst 200, duration 40s and temperature 20 °C. The fragmented DNA was used for metagenome preparation using the NEB Next Ultra II DNA library preparation kit. The DNA library was paired-end sequenced (2 x 150 bp) on a NovaSeq S4 system (Illumina, USA).

#### Oxford Nanopore sequencing

SQK-LSK114 sequencing libraries were prepared according to manufacturer recommendations with a minor custom modification to allow for native barcoding using kits EXP-NBD104 and EXP-NBD114 (Oxford Nanopore Technologies, Oxford, UK). Briefly; before initiating the SQK-LSK114 protocol, native barcodes were ligated onto end-prepped sample DNA (100-200 fmol) using NEB Blunt/TA ligase mastermix (New England Biolabs, USA). Approximately 10-20 fmol barcoded DNA library were loaded onto primed PromethION FLO-PRO114M (R10.4.1) flow cells and sequenced on the PromethION P2 Solo device running MinKNOW Release 22.07.3 (MinKNOW Core 5.3.0-rc3-p2solo).

#### Data preprocessing

Raw Illumina reads were filtered for PhiX using Usearch11^33^ and trimmed for adapters using cutadapt^34^ (v. 3.5). Forward and reverse read files were concatenated using a custom python script. Raw Oxford Nanopore fast5 files were basecalled and demultiplexed in Guppy v. 6.1.15 using the dna_r10.4.1_E8.2_400bps_sup algorithm. Adapters were removed in Porechop v. 0.2.4 using default settings. NanoStat v.1.4.0^35^ was used to assess quality parameters of the basecalled data. The adapter-trimmed data was then filtered in Filtlong v. 0.2.1 with – min_length set to 1500 bp and –min_mean_q set to 96 (q-score of 14).

#### Metagenome *de novo* assembly and binning

Metagenomes were assembled and binned using two independent pipelines in parallel. The resulting MAGs were lastly dereplicated in a single pool to produce the final 700 MAGs.

The first pipeline performed draft *de novo* co-assembly for metagenomes in six groups of samples/animals (combinations of control and treatment, corresponding to the seaweed supplementation, and a three-category methane variable representing low, medium and high emission levels) using Flye^36^ (v. 2.9.1-b1780) by setting the metagenome parameter (--meta). Draft metagenomes were first polished with Medaka^37^ (v. 1.7.1) using quality-filtered Oxford Nanopore R10.4.1 data, following further polishing with minimap2^38^ (v. 2.24-r1122) and racon^39^ (v. 1.5.0) using Illumina data covering the relevant metagenome sample trajectory. Each metagenome assembly was subjected to independent and automated genome binning using Metabat2^39^ (v. 2.15) and Vamb^40^ (v. 4.1.1). MAGs from each metagenome were subsequently dereplicated using dRep^41^ v. 3.2.2 setting minimum MAG length to 250000 bp (-l). All dereplicated MAGs from each metagenome assembly were finally pooled and dereplicated again (cross-dereplicated) with dRep.

The second pipeline accepted samples containing paired short-read and nanopore metagenomes. These were processed using a hybrid assembly approach, followed by MAG recovery through the Aviary^42^ (v. 0.5.7) pipeline (https://github.com/rhysnewell/aviary) using the recover workflow with default settings. The resulting assemblies were manually inspected using Bandage to identify and verify closed genomes.

The bins from both the first and second parallel pipelines were pooled, showcasing a total of 4,469 redundant recovered MAGs. Completeness and contamination rates were assessed with CheckM2^43^ (v. 1.0.1) using the lineage wf command. Only MAGs with >70% completeness and <10% contamination were retained for further analysis. To address potential multi-mapping issues during meta-omic relative abundance calculations, the genomes were dereplicated using a custom script. Pairwise Average Nucleotide Identity (ANI) values were calculated for all MAGs using Skani^44^. Genomes with >97% ANI and >50% alignment were clustered using complete linkage clustering. The highest-quality MAG within each cluster was selected as the representative genome. The quality score was calculated using the following metric: completeness - 5*contamination - 5*num_contigs/100 - 5*num_ambiguous_bases/100000, as described by Parks et al. (2020)^45^. This clustering process was iteratively repeated until no further clustering of representative MAGs was possible. This resulted in a nonredundant set of 700 MAGs.

### Genome-scale metabolic reconstruction and analysis

We built a genome-scale metabolic model (GEM) from each MAG using the automated metabolic reconstruction tool CarveMe^46^, gap filling for anaerobic growth on a complete medium (import allowed for all possible nutrients but not oxygen) to capture the metabolic environment of the rumen. We used the GEMs to convert relative archaeal and bacterial abundances from 16S rRNA sequencing and metagenomics data to metabolic reaction abundances. Annotating abundance data with the GTDB taxonomy^47^, we mapped the GEMs to the data by matching taxa on the genus level for the ASVs and the species level for the MAGs. We computed the frequency of each metabolic reaction for each ASV and MAG by taking the average of reaction presence (0 = reaction absent, 1 = reaction present) across all GEMs mapped on the genus level for the ASVs and by directly using reaction presence in GEMs for the MAGs. Then, we computed the abundance of each reaction in each sample by multiplying the reaction frequencies by the ASV or MAG abundances. We performed PCA separately for the ASV and MAG reaction abundances, standardizing features by removing the mean and scaling to unit variance.

### Rumen microbial genome database for metatranscriptomics and metaproteomics

For metatranscriptomic and metaproteomic data analyses, we built databases consisting of six parts representing different sources and taxonomic domains:

A. 700 MAGs assembled from our digesta samples, representing both archaea and bacteria.
B. *Bos taurus* host genome ARS-UCD1.3^48^ GCF_002263795.2 (NCBI Bioproject PRJNA391427).
C. *Entodinium caudatum* genome^49^ (NCBI Bioproject PRJNA380643).
D. 52 protozoal SAGs^2^ (NCBI Bioproject PRJNA777442).
E. 9 fungal genomes from phylum Neocallimastigomycota^10^.
F. 14 bacterial genomes of genus *Campylobacter*^50^.

This total rumen microbial genome database consisted of 4.2 M proteins with an average length of 426.8 amino acids totalling 1.8 G amino acid letters.

#### Annotation of genomes and characterization of proteins

The different parts of the rumen microbial genome database (A-F) were annotated using several tools. For the 700 recovered MAGs (rumen microbial genome database part A) and 14 *Campylobacter* spp. genomes (F), Prokka^51^ (v. 1.14.6) was used for annotation and to translate the coding sequences. CheckM2^52^ (v. 1.0.2) was used for assessment of completeness and contamination. The remaining database parts (B–E) were downloaded as amino acid sequences. Translated genes of the complete rumen genome database (A–F) were characterized functionally using eggnog-mapper^53^ (v. 2.1.12), resulting in the identification of PFAM^54^, CAZy^55^, and KEGG^56^ orthologs. The eggnog-mapper results were predominantly used for interpretation of the (meta)proteomic analysis. Pathway enrichment analysis was calculated using the KEGG orthologs and KEGG pathway database^57^ (downloaded on 2023-08-28) via clusterProfiler^58^ (v. 4.10.0). Taxonomic identification of MAGs were done with GTDB-tk^47^ (v. 2.4.0) using database r214. The genomic characterization tools mentioned above were run via CompareM2^59^ (v. 2.11.1). For screening of metabolic capacities, DRAM^60^ (v. 1.4) was used on the translated amino acid sequences of the complete rumen microbial genome database with the following parameters: DRAM.py annotate_genes --use_uniref --threads 64. The DRAM results were predominantly used for interpretation of the (meta)transcriptomic analysis. CoverM^61^ (v. 0.6.1) was used to calculate read coverage and estimate relative abundances of bacterial and archaeal MAGs (A).

### Meta- and host transcriptomics

RNA extraction and sequencing for rumen digesta, wall and liver samples, as well as bioinformatic analyses for rumen wall and liver sequence data, were performed at DNASense ApS (Aalborg, Denmark).

#### RNA extraction

RNA extraction for rumen digesta, rumen wall and liver samples was performed with the standard protocol for RNeasy PowerMicrobiome Kit (Qiagen, Germany) with minor modifications: custom reagent volumes were used, PM4 buffer was replaced with 70 % ethanol in initial extraction mix, and bead beating was performed at 6 m/s for 4×40s. Gel electrophoresis using Tapestation 2200 and RNA screentape (Agilent, USA) was used to validate product integrity and purity of RNA extracts. RNA concentrations were measured using Qubit RNA HS/BR Assay kit (Thermo Fisher Scientific, USA). The extracted RNA was treated with the TURBO DNAfree (Thermo Fisher Scientific, USA) to ensure removal of all DNA in the samples. Afterwards the RNA was quality controlled using RNA screentape (Agilent, USA) and Qubit RNA HS/BR Assay kit (Thermo Fisher Scientific, USA).

#### Sequencing library preparation

RNA extracts were rRNA depleted using the Ribo-Zero Plus rRNA Depletion Kit (Illumina, USA), and residual DNA from RNA extraction was removed using the DNase MAX kit (MoBio Laboratories Inc.). The samples were purified using the standard protocol for CleanPCR SPRI beads (CleanNA, NL) and further prepared for sequencing using the NEBNext Ultra II Directional RNA library preparation kit (New England Biolabs). Library concentrations were measured using Qubit HS DNA assay and library DNA size estimated using TapeStation with D1000 ScreenTape. The samples were pooled in equimolar concentrations and sequenced (2 x 150 bp, PE) on a Novaseq platform (Illumina, USA). All kits were used as per the manufacturer’s instructions with minor modifications.

#### Host transcriptome mapping

Forward and reverse sequencing cDNA reads were quality-filtered and trimmed for Illumina adapters using Cutadapt v. 3.7^62^ used in paired-end mode. For liver and rumen wall data, the reads were subsequently mapped against the Bos Taurus Genome Reference ARS-UCD1.3 (Genbank assembly accession GCA_002263795.3). The genome and its associated gene transfer format file (GTF) were downloaded and indexed using STAR^63^ (v. 2.7.10a), setting a sjdbOverhang of 149 bp. Adapter-trimmed sample reads were mapped against the indexed genome of ARS-UCD1.3 using STAR (v. 2.7.10a) in paired-end mode, with the option - outReadsUnmapped Fastx enabled. Alignments were ported to coordinate-sorted BAM files, and FeatureCounts (v. 2.0.1) of the SubRead package^64^ was used to quantify CDS mappings as counts. Where nothing else is stated, the default settings were used for all bioinformatic tools.

#### Rumen wall metatranscriptome mapping

For rumen wall samples, the forward and reverse cDNA reads that did not map against the *Bos taurus* genome were bioinformatically depleted for rRNA using Ribodetector v. 0.2.7^65^ and then mapped against the rumen MAGs. Prior to mapping, the concatenated MAGs were indexed using STAR^66^ (v. 2.7.10a). The rRNA-depleted and quality filtered DNA reads were mapped against the MAGs with STAR, setting alignIntronMax to 1. All alignments were ported to coordinate-sorted BAM files.

#### Rumen content metatranscriptomics

Rumen content data were mapped against the Bos taurus genome (Genome Reference ARS-UCD1.3) using minimap2 v 2.2. All non-paired mapped reads were retrieved using samtools v 1.17^67^ with the following parameters samtools fastq -f 12 -F 256 -c 7 -1 read1.fq.gz -2 read2.fq.gz. rRNA reads present in the samples were bioinformatically removed using SortMeRNA v 4.3.6^68^ with the following SILVA databases: silva-bac-16s-id90, silva-arc-16s-id95, silva-bac-23s-id98, silva-arc-23s-id98, silva-euk-18s-id95 and silva-euk-28s-id98, and the parameters –out2–paired_out –fastx–thread 64. These reads were used to quantify the expression of coding sequences (CDS) encoded in the rumen microbial genome database using Kallisto^69^ (v. 0.50.0). The resulting ‘raw-counts’ tables were gathered into a single table using the Bioconductor tximport^70^ (v. 1.26.1) library in R 4.2.2.

### Meta- and host proteomics

Proteomic and metaproteomic measurements and all bioinformatic analyses for rumen digesta, wall and liver samples were performed at the Norwegian University of Life Sciences (NMBU; Ås, Norway).

#### Protein extraction and digestion

Protein extraction was performed following a previously published protocol^1^. Briefly, for rumen samples we used 300 μL of fluid for downstream processing; for liver samples we used ∼300μL of finely chopped/liquified liver (with sterile scalpel); finally for rumen wall samples we used sterile tweezers and scalpel to carefully remove the wall papillae from the remainder of the tissue and finely chop them into liquified mass (∼300μL). Each sample was combined with 150 μL lysis buffer (30 mM DTT, 150 mM Tris-HCl (pH = 8), 0.3% Triton X-100, 12% SDS) and 4 mm glass beads (≤160 μm), then vortexed and rested on ice for 30 mins. Sample lysis was performed with a FastPrep-24 Classic Grinder (MP Biomedical, Ohio, USA) for 3 × 60 s at 4.0 m/s^71^, followed by centrifugation at 16,000 × g for 15 min at 4 °C. Lysate was removed and its absorbance measured at A750 on BioTek Synergy H4 Hybrid Microplate Reader (Thermo Fisher Scientific Inc., Massachusetts, USA). 40–50 μg of protein was prepared in SDS-buffer, heated in a water bath for 5 min at 99 °C, and analyzed by SDS-PAGE with Any-kD Mini-PROTEAN TGX Stain-Free gels (Bio-Rad, California, USA) in a 2 minute run for sample clean-up, before staining with Coomassie Blue R-250. Visible bands were excised and divided into 1 mm^2^ pieces before reduction, alkylation and trypsin digestion. Peptides were concentrated and eluted using C18 ZipTips (Merck Millipore, Darmstadt, Germany) following manufacturer’s instructions.

#### Mass spectrometry

Peptide samples were analyzed by coupling a nano UPLC (nanoElute, Bruker) to a trapped ion mobility spectrometry/quadrupole time of flight mass spectrometer (timsTOF Pro, Bruker). Peptides were separated with a PepSep Reprosil C18 reverse-phase (1.5 µm, 100Å) 25 cm X 75 μm analytical column coupled to a ZDV Sprayer (Bruker Daltonics, Bremen, Germany). Column temperature was kept at 50°C using the integrated oven. Equilibration of the column was performed before the samples were loaded (equilibration pressure 800 bar). The flow rate was set to 300 nl/min and the samples separated using a solvent gradient from 5 % to 25 % solvent B over 70 minutes, and to 37 % over 9 minutes. The solvent composition was then increased to 95 % solvent B over 10 min and maintained at that level for an additional 10 min. In total, a run time of 99 min was used for the separation of the peptides. Solvent A consisted of 0.1 % (v/v) formic acid in milliQ water, while solvent B consisted of 0.1 % (v/v) formic acid in acetonitrile.

The timsTOF Pro was run in positive ion data dependent acquisition PASEF mode with the control software Compass Hystar version 5.1.8.1 and timsControl version 1.1.19 68. The acquisition mass range was set to 100 – 1700 m/z. The TIMS settings were: 1/K0 Start 0.85 V⋅s/cm2 and 1/K0 End 1.4 V⋅s/cm2, Ramp time 100 ms, Ramp rate 9.42 Hz, and Duty cycle 100 %. Capillary Voltage was set at 1400 V, Dry Gas at 3.0 l/min, and Dry Temp at 180 ℃. The MS/MS settings were the following: number of PASEF ramps 10, total cycle time 0.53 sec, charge range 0-5, Scheduling Target Intensity 20000, Intensity Threshold 2500, active exclusion release after 0.4 min, and CID collision energy ranging from 27-45 eV.

#### Data analysis

The raw spectra were analyzed using mspipeline1^72^ (v. 2.0.0) based on FragPipe^73^ (v. 19.1). Using Philosopher^74^ (v. 4.8.1), MSFragger^75^ (v. 3.7) and IonQuant (v. 1.8.10). Spectra were analyzed slicing the rumen microbial genome database into 16 parts using msfragger.misc.slice-db=12. Mass calibration was disabled with msfragger.calibrate_mass=0. The maximum length of peptides to be generated during in-silico digestion was 35 with msfragger.digest_max_length=35. Allowed number of missed cleavages 1 and 2 was set to 1 with msfragger.allowed_missed_cleavage_{1,2}=1. Otherwise, default settings were used. The processing was performed on an AMD x86-64 “Threadripper Pro” 5995WX 64 cores, 8 memory channels, 512GiB DDR4 3200MHz ECC (8x 64 GiB) and 4 2TB SSDs in raid0.

Proteomic intensities were log2-transformed prior to any statistical analysis. Genes in the proteomic database were annotated using eggnog e-mapper (v. 2.1.12) using CompareM2 (v. 2.11.1). Missing values were imputed using missRanger^76^ (v. 2.6.0).

### Untargeted metabolomics

Untargeted metabolomic analyses for rumen digesta, rumen wall, and liver samples were carried out by MS-Omics Aps (Vedbæk, Denmark). Compound identification was performed at four levels: Level 1: identification by retention times (compared against in-house authentic standards), accurate mass, and MS/MS spectra; Level 2a: identification by retention times (compared against in-house authentic standards), and accurate mass; Level 2b: identification by accurate mass, and MS/MS spectra; Level 3: identification by accurate mass alone. A deviation of 3 ppm was accepted for accurate mass identification.

#### Sample extraction

Rumen digesta samples were vortexed and an aliquot (100 µl) transferred to a spin filter (0.22µm). The aliquot was diluted with water (100 µl) and filtered by centrifugation (7000 rpm, 2 x 5 min, 4°C). Filtered extracts were diluted 10 times in mobile phase eluent A and fortified with stable isotope labeled standards before analysis.

Rumen wall and liver samples were mixed with ceramic beads and precooled methanol/water (1:2) fortified with stable isotope labeled standards. The samples were then placed in a pre-cooled (-20°C) bead beater and homogenized (4 x 30 sec., 30 Hz) followed by ultrasonication (5 min). After centrifugation (18000 RCF, 5 min, 4°C), the supernatant of each tube was collected. The sample pellets were re-extracted as described above. The two extract supernatants were pooled and passed through a phosphor removal cartridge (Phree, Phenomenex). A precise aliquot of the extract was evaporated to dryness under a gentle stream of nitrogen, before reconstitution with 10% Eluent B in Eluent A.

#### LC-MS method

All samples were analyzed using a Thermo Scientific Vanquish LC coupled to an Orbitrap Exploris 240 MS instrument (Thermo Fisher Scientific). An electrospray ionization interface was used as the ionization source. Analysis was performed in positive and negative ionization mode under polarity switching. Ultra-performance liquid chromatography was performed using a slightly modified version of a published protocol^77^. Peak areas were extracted using Compound Discoverer (Thermo Scientific) version 3.2 (digesta) or 3.3 (liver and wall). For the wall and liver samples, probabilistic quotient normalization was applied prior to further analyses, to decrease concentration effects.

### Volatile fatty acid quantification

Rumen digesta samples were thawed on ice and centrifuged when still cold. 450 µL of each sample was transferred to a new tube and 50 µL of a 50% formic acid solution was added to reach a 5 % concentration of formic acid. Samples were then centrifuged again and 400 µL of the supernatant was transferred to a GC-vial, with 1000 µL of an internal standard solution added. Volatile fatty acids were separated using gas chromatography (Trace 1300 GC with autosampler, Thermo Scientific) with a Stabilwax-DA column (30m, 0.52 mm ID, 0.25 µm, Restek).

### Microarray polymer profiling

Microarray polymer profiling (MAPP) entails the printing of extracted glycans as high-density microarrays which are then probed with monoclonal antibodies with specificities for different glycan epitopes. The output from MAPP provides insight into the relative abundance of epitopes across the sample set.

Alcohol insoluble residues (AIR) were prepared from each rumen digesta sample (n=24) as follows: samples were homogenized to a fine powder using a tissue lyser (Qiagen). Approximately five volumes of 70% ethanol were added, the samples vortexed for 10 minutes then centrifuged at 2,700 g for 10 minutes and the supernatants discarded, This step was repeated. Approximately five volumes of 1:1 methanol and chloroform were added to the pellet and the samples were again vortexed and centrifuged as previously. Finally, approximately five volumes of acetone were added and the same vortexing and centrifugation steps performed. The resulting pellets were AIR.

To extract glycans, 300 μL of 50 mM diamino-cyclo-hexane-tetra acetic acid, pH 7.5, were added to 10 mg AIR. After agitation in a tissue lyser (27 s-1 for 2 minutes and 10 s-1 2 hours), samples were centrifuged at 2,700 g for 10 minutes. The supernatant was removed, 300 μL 4M NaOH with 1% v/v NaBH4 added to the pellet and the agitation and centrifugation steps repeated. The resultant NaOH extraction supernatants were diluted sequentially (1/2,1/5,1/5,1/5) in microarray printing buffer (55.2% glycerol, 44% water and 0.8% Triton X-100), and the four dilutions were printed in quadruplet onto nitrocellulose membranes using a non-contact microarray robot (Arrayjet, Roslin). Thus, every replicate was represented by a 16-spot subarray (four concentrations and four printing replicates). Arrays were probed with monoclonal antibodies, scanned, uploaded into microarray analysis software (Array Pro Analyzer 6.3, Media Cybernetics) and mean spot signals from each sub array calculated.

### Statistics and data visualization

Unless otherwise specified, statistical analyses and visualizations were performed in the R statistical programming language^78^ (v. 4.3.2). The knitr^79^ package (v. 1.45) was used for reporting, renv^80^ (v. 1.0.7) for package management, ggplot2^81^ (v. 3.5.1) for visualizations, cowplot^82^ (v. 1.1.3) for composing multipanel figure layouts, and ComplexHeatmap^83^ (v. 2.15.4) for heatmaps.

16S rRNA gene ASV data was managed with phyloseq^84^ (v. 1.46.0), which was also used to calculate alpha diversity indices. Rumen community types (RCTs) were defined using the ASVs data and Dirichlet Multinomial Mixtures^85^ clustering implemented with mia^86^ (v. 1.10.0), selecting the optimal number of clusters based on the Laplace method. Only ASVs that were present in at least half of all slaughter timepoint samples (n = 35) were used for this analysis.

All beta diversity comparisons for ASV counts and MAG relative abundances were performed using vegan^87^ (v. 2.6-6), with robust Aitchison distances statistically compared with PERMANOVA (adonis2; 9999 permutations), and visualized with PCoA (package ape^88^, v. 5.8). Principal Component Analysis (PCA) for all other omic data types was run with the “prcomp” function. For (meta)transcriptomic data (variance stabilizing transformed (VST) counts) and (meta)proteomic data (log2 transformed LFQ intensities with imputed missing values), the 1000 features with the highest variance were selected for PCA. For untargeted metabolomic data, where the number of features was orders of magnitude lower, all features were used, except for rumen digesta, where features with annotation level 3 were excluded.

Where statistical testing was done between two categorical variables, Fisher’s exact tests were used. Continuous variables were compared either with t-tests (KPTs and other animal-related metrics, Principal Component scores), or Wilcoxon rank sum tests (alpha diversity indices, metagenomic relative abundances, proteomic LFQ intensities, MAPP intensities, normalized intensities of metabolites from untargeted measurements, and molar percentages of volatile fatty acids) with multiple comparison correction using the “fdr” option of the “p.adjust” function. Differential abundance comparisons for count data (ASVs and meta- and host transcriptomics) were run with DESeq2^89^ (v. 1.42.1), with default parameters for transcriptomic data, and the “sfType” parameter changed to "poscounts" for ASV data.

#### Network analysis (WGCNA)

Correlation-network based analysis was applied on the proteomic and metaproteomic samples to group co-expressed proteins into clusters. Weighted gene co-expression network analysis^21^ (WGCNA) (v. 1.73) was applied on data that included imputed missing values to construct clusters independently in the digesta, rumen wall epithelium, and liver samples. These clusters were then correlated via their eigengenes across samples to obtain host-microbiome boundary-links.

## Acknowledgements

We gratefully acknowledge the financial support of the Novo Nordisk Foundation under 0054575-SuPAcow. This project has received funding from the European Union’s Horizon 2020 Research and Innovation programme under grant agreement number No.101000213. PBP also acknowledges support from the Australian Research Council (Future Fellowship: FT230100560). The authors acknowledge the Orion High Performance Computing Center at the Norwegian University of Life Sciences and Sigma2—the National Infrastructure for High Performance Computing and Data Storage in Norway for providing computational resources that have contributed to computations reported in this paper. We also acknowledge Elixir Norway, supported by the Research Council of Norway’s (NFR) grant 322392, for the bioinformatics and data management support received for this paper. The authors further acknowledge financial support from the Scottish Government (RESAS Division) and Biotechnology and Biological Sciences Research Council (BBSRC BB/S006567). We also thank the staff of the SRUC Beef Research Centre for their excellent technical support. BioRender.com is acknowledged for providing the platform used to create Fig. 1 and 5.

## Extended Data Figures

**Extended Data Fig 1.**
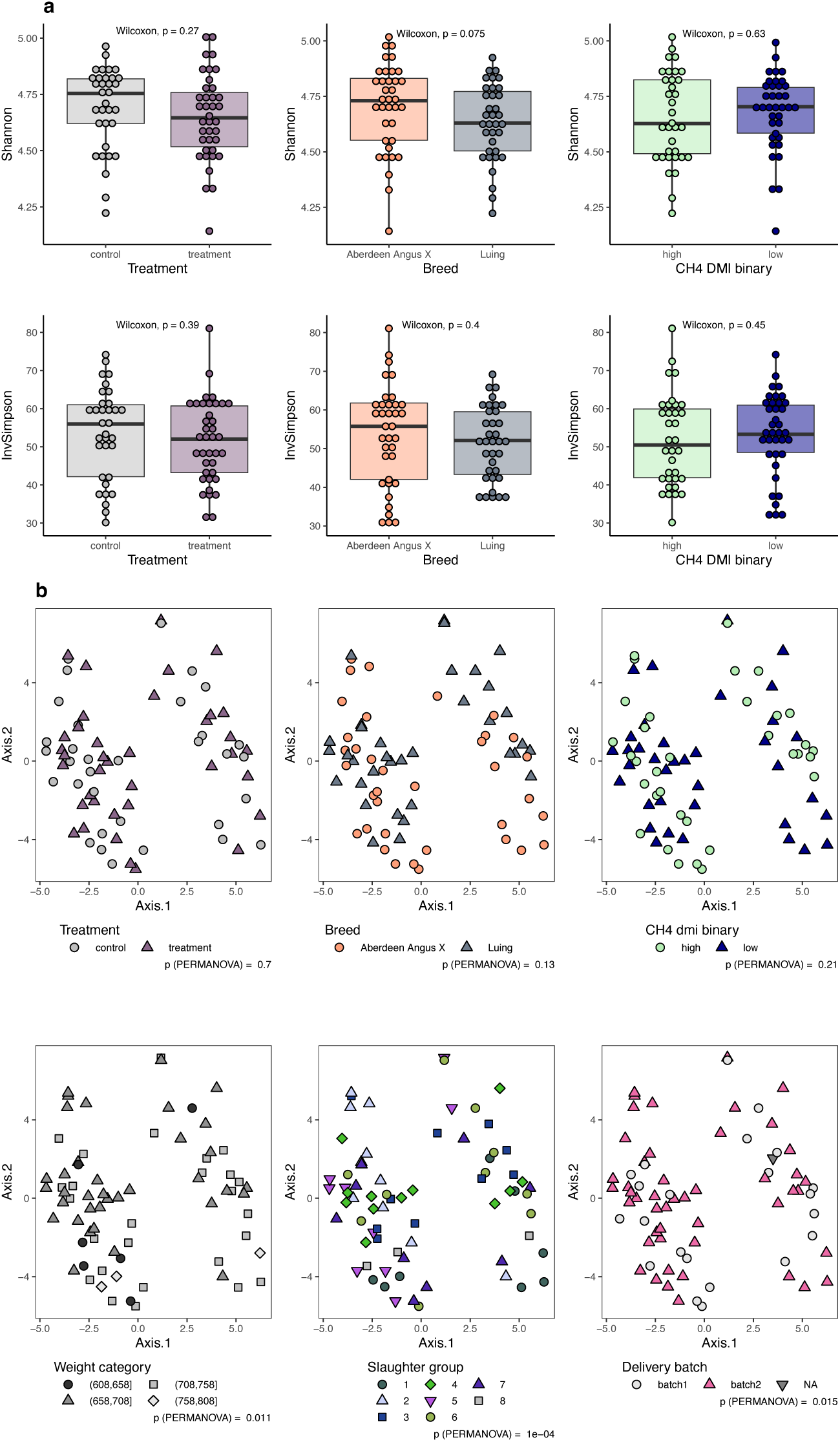
**16S rRNA gene amplicon sequence data analysis results for 71 animals**. **a**. Alpha diversity, showing Shannon and inverse Simpson indices for treatment (short-term seaweed supplementation), breed (Aberdeen Angus X vs Luing) and methane emission level (binary categorical variable based on median CH_4_ g/kg DMI). Box hinges represent the 1st and 3rd quartiles; whiskers range from hinge to the highest and lowest values that are within 1.5*IQR of the hinge. **b**. Beta diversity visualized with Principal Coordinates Analysis of robust Aitchison distances and statistically compared with PERMANOVA (adonis2), showing the three main grouping variables (treatment, breed and methane emission category) as well as the three tested variables with the lowest *p*-value (liveweight, slaughter group and delivery batch). All *p*-values are shown without multiple comparison correction.

**Extended Data Fig 2.**
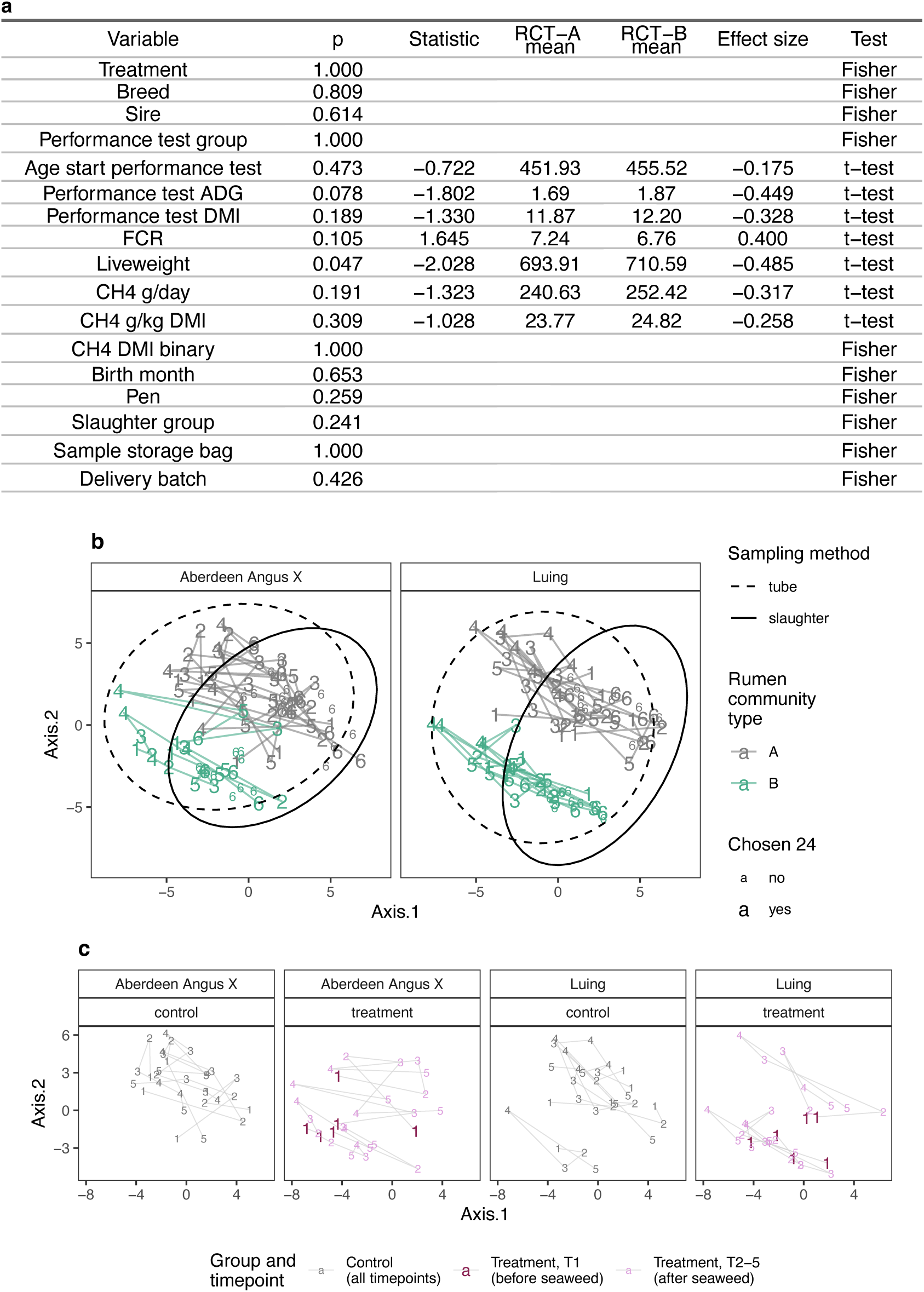
Comparisons of rumen community types and technical and animal-related variables. **a.** Statistical comparisons of variables against the two community types, with t-tests for numerical variables and Fisher’s exact tests for categorical variables. All p-values are shown without multiple comparison correction. **b.** Principal Coordinates Analysis (PCoA) with robust Aitchison distances of rumen digesta 16S rRNA gene amplicon sequence data, showing samples from six time points that spanned the length of the animal trial. Numbers correspond to timepoint, colors to rumen microbial community type, and the 24 animals chosen for deeper analysis are indicated with larger symbols. Samples from the same animal are connected with lines in sequential order. Ellipses indicate 95% confidence levels for sample types: tube sampling (timepoints 1-5; dashed line) or post-slaughter sampling (timepoint 6, continuous line). **c.** The same PCoA plot as presented in **b**, with lines connecting samples from the same animal, but showing only tube-collected samples, grouped by treatment group (control or seaweed supplementation) in addition to breed, and indicating the before and after seaweed timepoints.

**Extended Data Fig 3.**
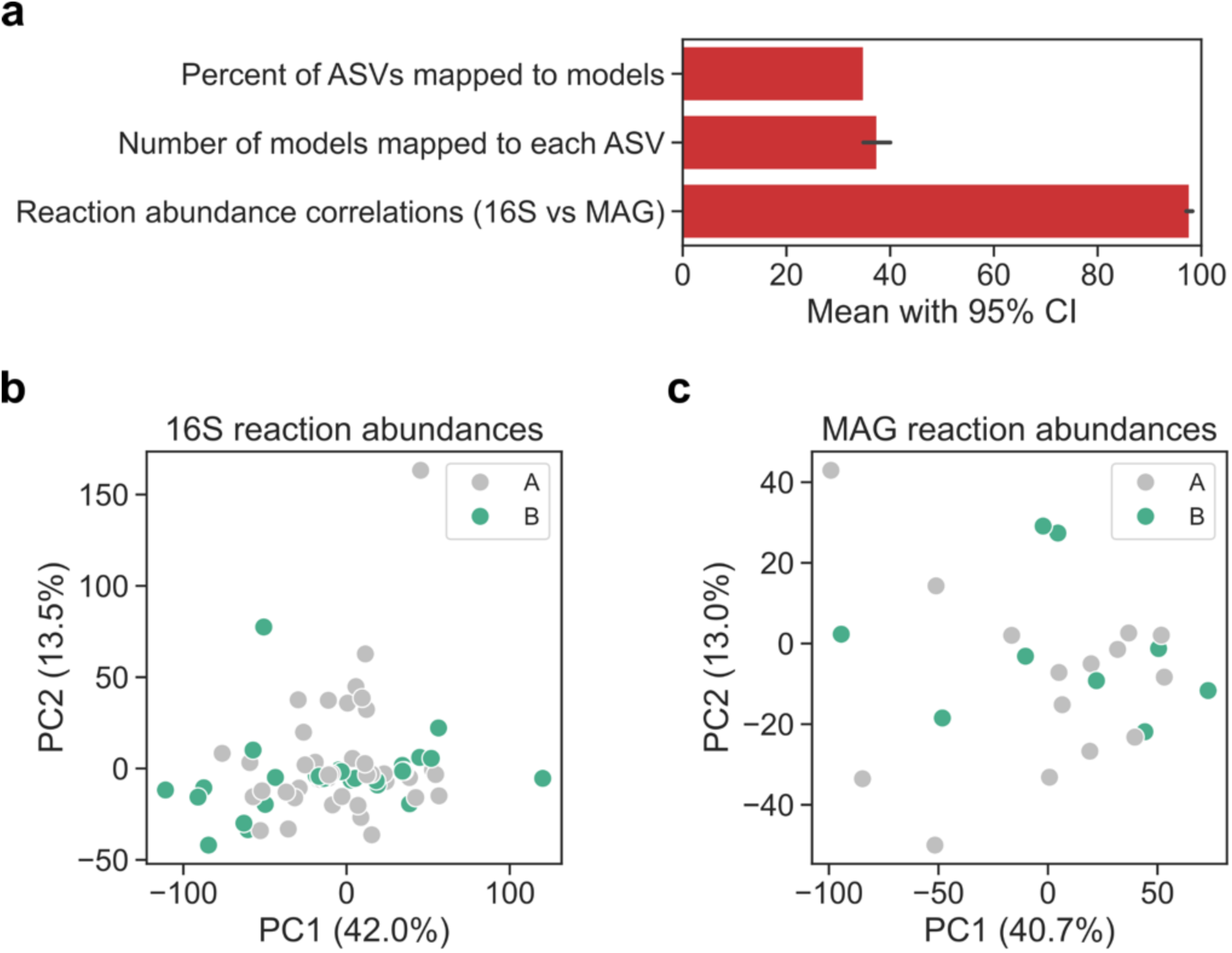
Metabolic reaction abundance analysis. **a.** Mapping of ASVs from 16S rRNA gene sequencing data to genome-scale metabolic models of archaea and bacteria built from 700 MAGs. From top to bottom: ASVs taxonomically mapped at the genus level to at least one model (%), models taxonomically mapped at the genus level for each ASV, and Pearson correlation (%) between metabolic reaction abundances calculated from ASV and MAG abundances for each sampled animal. Means are shown with 95% confidence intervals from bootstrapping with 1000 samples **b.** PCA score plot of metabolic reaction abundances calculated from ASV abundances. **c.** PCA score plot of metabolic reaction abundances calculated from MAG abundances. In each score plot, PC1 and PC2 are shown with their explained variance (%). Each dot is a sampled animal and color indicates RCT.

**Extended Data Fig 4.**
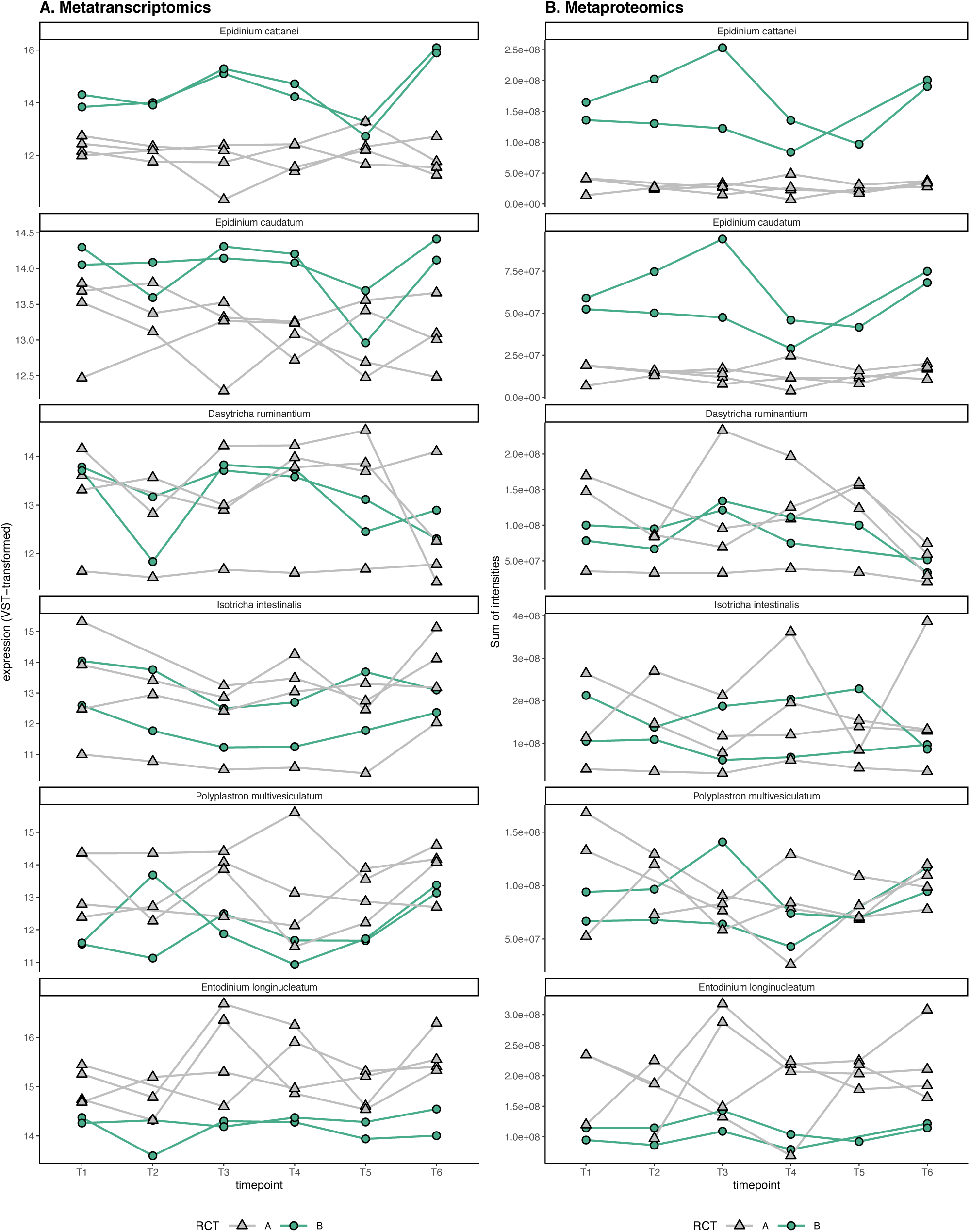
Abundances of select protozoal species in rumen digesta samples over time. Sampling for timepoints T1-T5 was performed through a stomach tube, while T6 was collected post-slaughter. Lines connect samples from the same animal (with a total of 6 animals with time series data). Shape and color correspond to rumen community type (RCT-A or -B). **a**. VST-normalized sums of counts per ciliate species for metatranscriptomic data. **b**. Sums of LFQ intensities per ciliate species for metaproteomic data, excluding samples with < 1000 ciliate protein groups detected. All plots have one missing data point (T2 from an animal in the RCT-A group) for which the tube sampling was not successful.

**Extended Data Fig 5.**
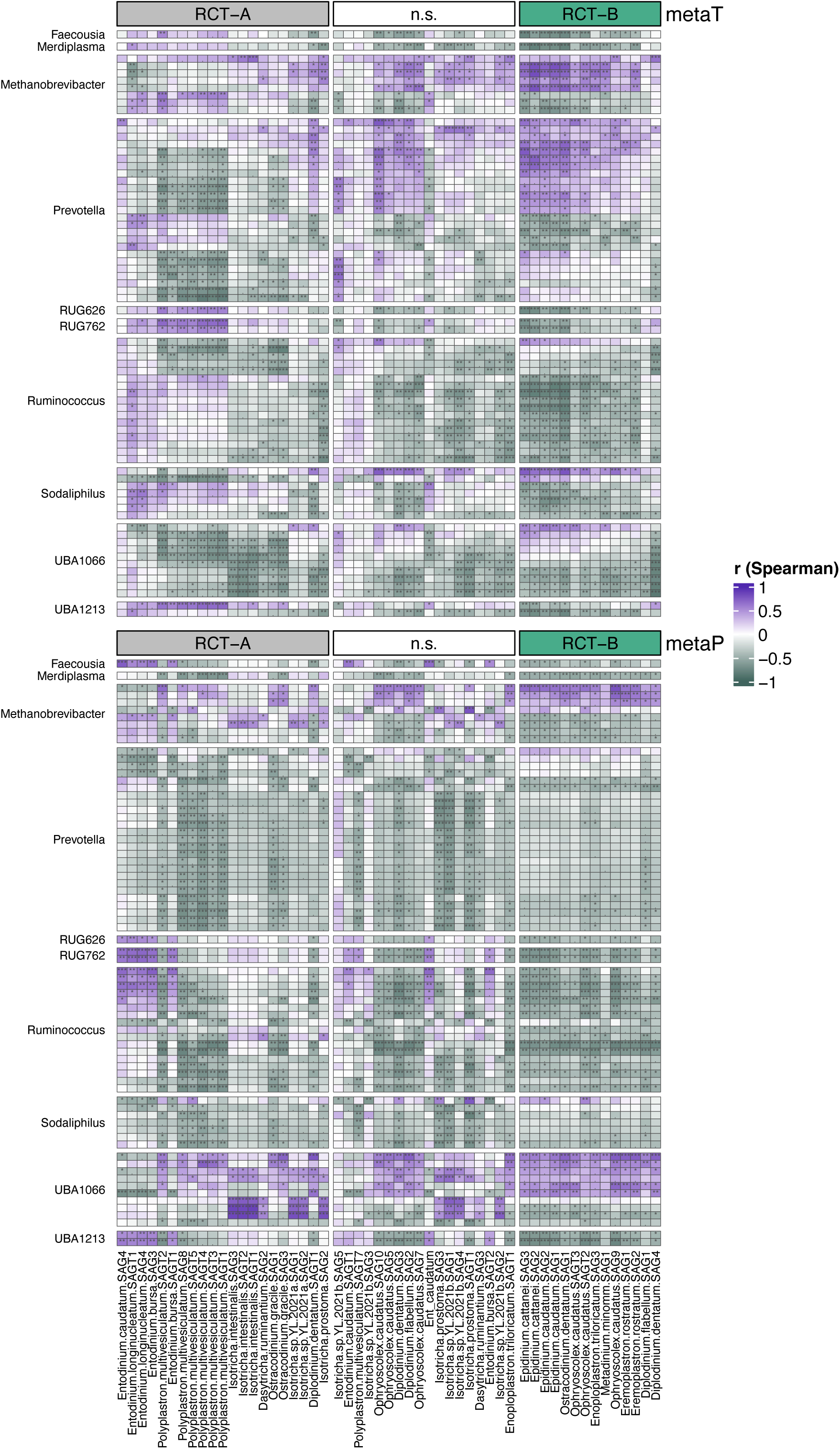
Heatmap of Spearman correlations between archaea and bacteria of interest and ciliates in metatranscriptomic and metaproteomic data. Rows correspond to individual archaeal and bacterial MAGs, and are labeled according to the genus classification of the MAGs. Columns correspond to individual ciliate SAGs (and “Ent_caudatum”, representing the first published genome available for *Entodinium caudatum*), ordered according to their differential abundance between the two rumen community types (RCT-A and -B) in either metatranscriptomic (metaT) or metaproteomic (metaP) data (if both or either suggest enrichment in one RCT with adjusted *p* < 0.1, the SAG is listed under that RCT; if there is a discrepancy between the two omics, the SAG is counted as not significant (n.s.)). The plot shows those MAGs that were among the genera with highest loadings in Fig 2g, and are significantly correlated (p < 0.05) with at least 10 SAGs in both metatranscriptomic and metaproteomic data; the same MAGs are shown in both the metaT and the metaP parts of the heatmap. Stars reflect uncorrected *p-*values as follows: *** : p < 0.001, ** p < 0.01, * : p < 0.05, . : 0.1 > p > 0.05.

**Extended Data Fig 6.**
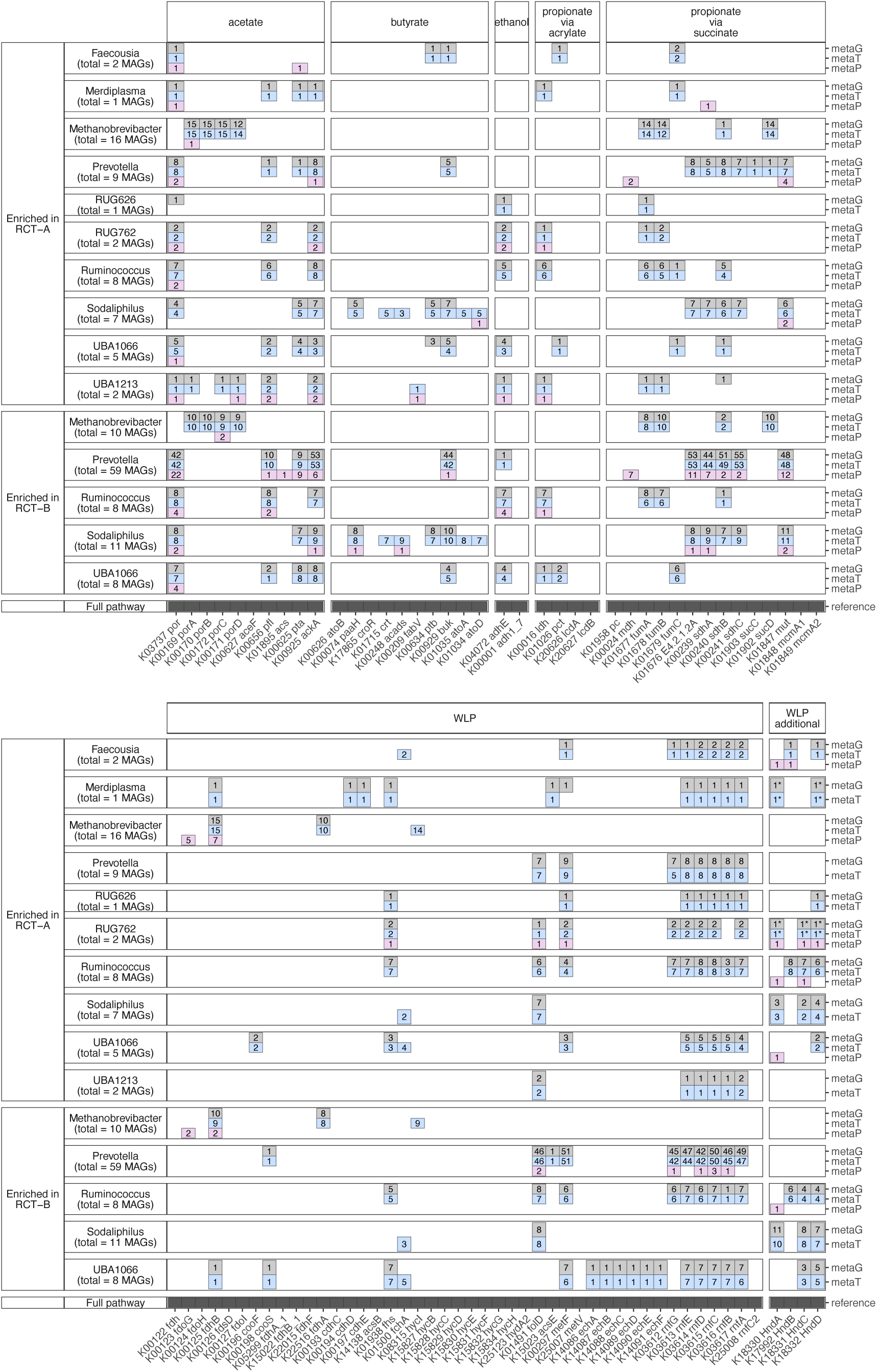
Summary of KOs in pathways of interest detected in MAGs. Numbers reflect how many MAGs out of the genera of interest had the indicated KOs present in each omic layer. The “1*” label for RUG762 and Merdiplasma Hnd KOs indicates that due to different annotation approaches, the KOs were not initially detected in metaG/metaT, but their presence is strongly indicated based on manual sequence exploration using the HydDB (Søndergaard et al. 2016) and Conserved Domain Database (Wang et al. 2023).

**Extended Data Fig 7.**
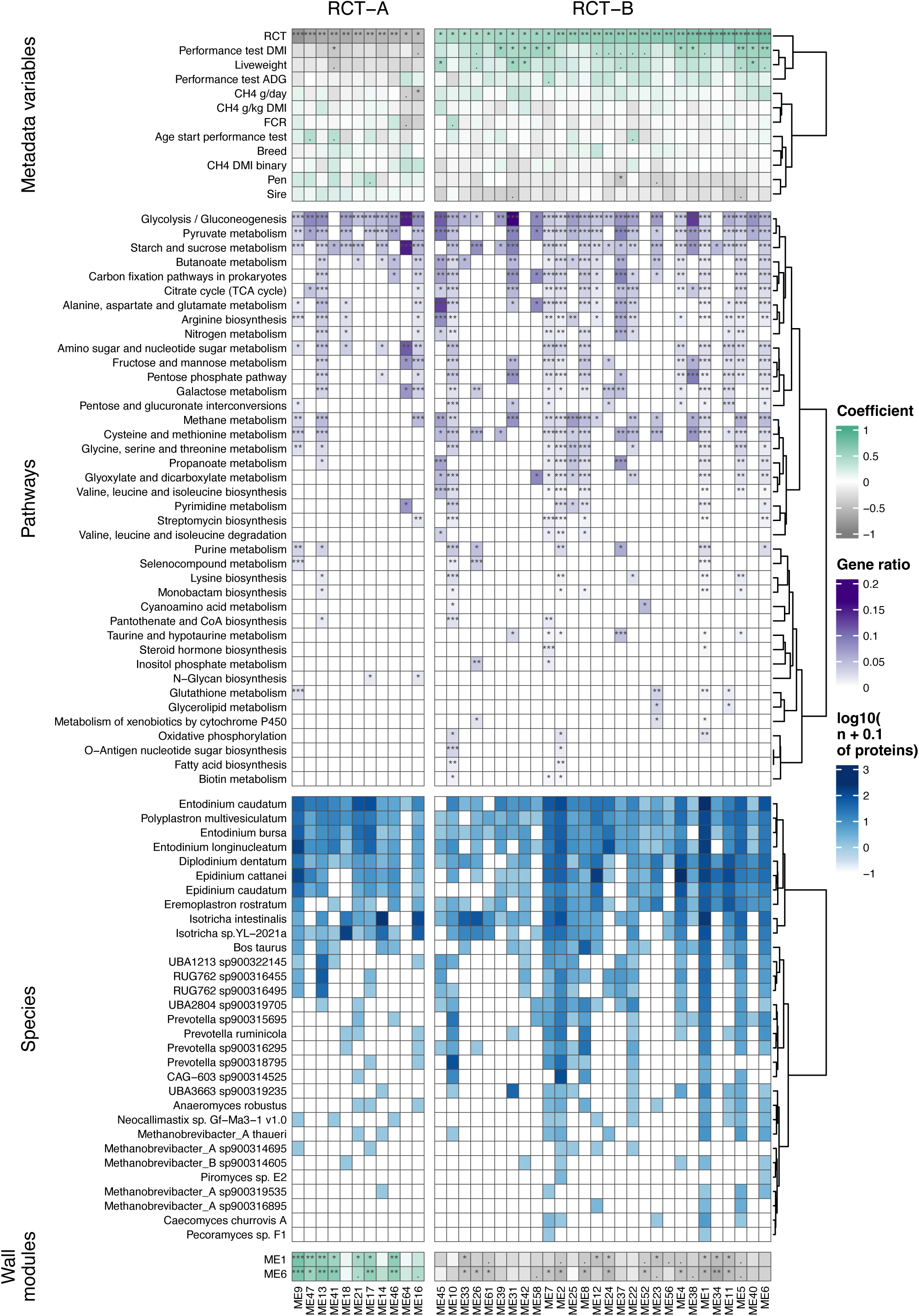
**Heatmap of weighted gene co-expression network analysis results of rumen digesta metaproteomics**. Showing the 39 modules out of a total of 65 that had p < 0.05 for Pearson correlations with rumen community types (RCT). “Metadata variables” rows show Pearson correlations between modules and animal metrics. “Pathways” rows show those KEGG pathways of class "09100 Metabolism" that were significantly enriched using the hypergeometric test in more than one RCT-correlated module. “Species” rows show the numbers of protein groups per species assigned to the modules, including the top ten taxa with the most protein groups in the RCT-correlated modules for bacteria and protozoa each, and the top five for archaea and fungi. The “Wall modules” rows indicate biweight midcorrelation between digesta and wall modules.

**Extended Data Fig 8.**
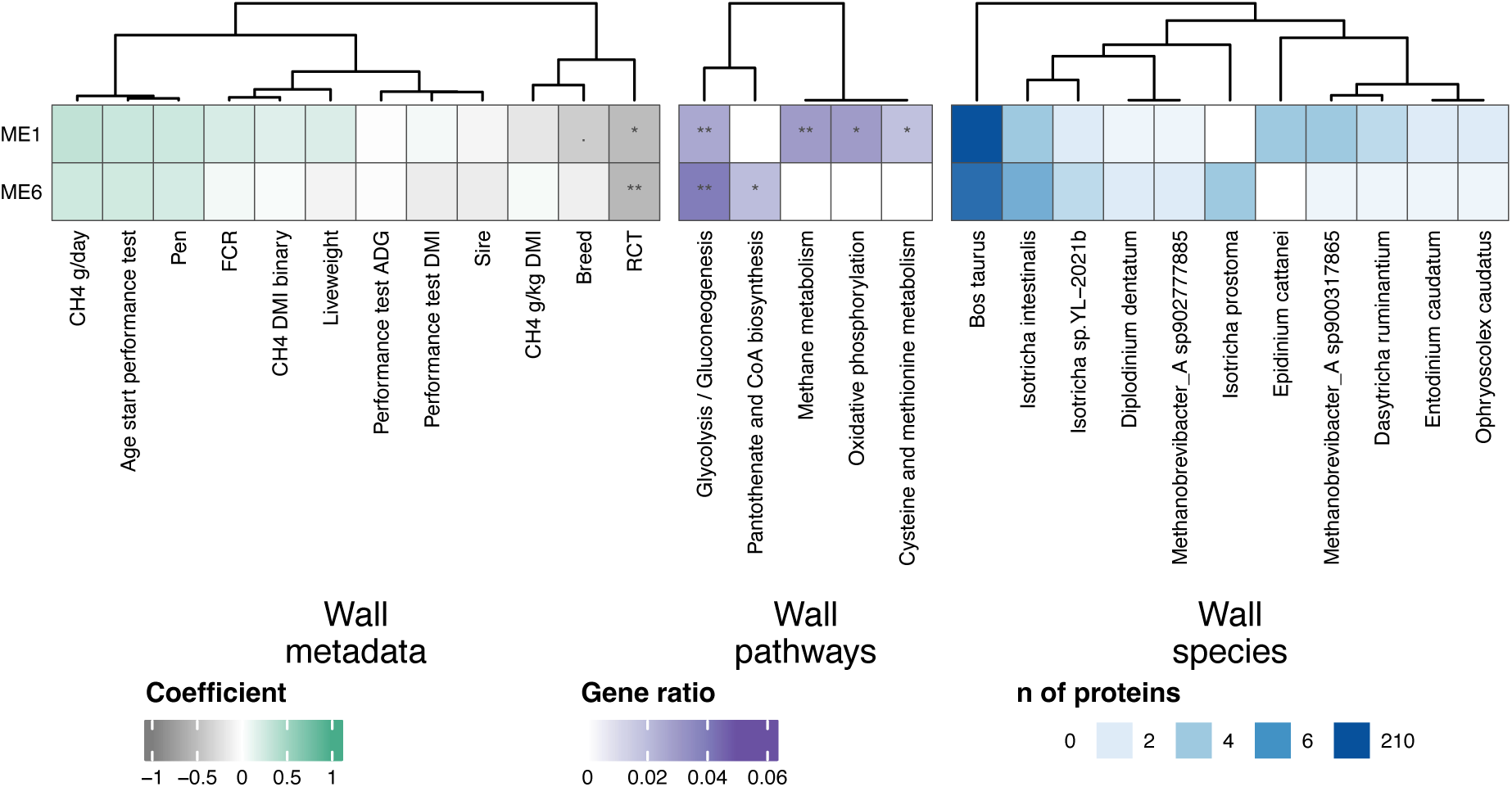
**Heatmap of weighted gene co-expression network analysis results of rumen wall metaproteomics**. Showing the 2 modules out of a total of 19 that had p < 0.05 for Pearson correlations with rumen community types (RCT). “Wall metadata” columns show Pearson correlations between modules and animal metrics. “Wall pathways” columns show those KEGG pathways of class "09100 Metabolism" that were significantly enriched using the hypergeometric test in more than one RCT-correlated module. “Wall species” columns show the numbers of protein groups per species assigned to the modules, including all taxa with more than 2 protein groups detected.

